# *Arabidopsis* YEATS domain proteins facilitate DNA double strand break repair via homologous recombination pathways

**DOI:** 10.1101/2025.10.11.681831

**Authors:** Neeraja Vegesna, Clara Bourbousse, Yasaman Jami-Alahmadi, Ana Marie S. Palanca, James A. Wohlschlegel, Julie A. Law

## Abstract

DNA double-strand break (DSB) repair is required to maintain genome integrity and organismal fitness. This process occurs within chromatin and although accurate repair requires extensive changes in chromatin structure and composition, chromatin effectors facilitating DSB repair remain largely unknown in plants. Using reverse genetics, we identified two chromatin readers, YAF9A and YAF9B, as new factors required for DSB repair in *Arabidopsis*. While only *YAF9B* is induced by DNA damage, both YAF9A and YAF9B are required for DSB repair, including via homologous recombination (HR)-like pathways. Mechanistically, we found YAF9A and YAF9B denote distinct remodeling complexes: While YAF9A associates with both NuA4 and SWR1, YAF9B only associates with NuA4, defining a DNA-damage-specific version of this remodeling complex and revealing the first link between NuA4 and DSB repair in plants. We also uncovered direct roles for YAF9A and YAF9B in DNA repair, rather than indirect roles in gene regulation, as *yaf9* mutants showed normal transcriptional responses to DNA damage. Finally, we demonstrate that the YAF9B reader domain is required for DSB repair. These results link YAF9A, YAF9B, and their respective remodeling complexes, to DSB repair in a histone reader-dependent manner, expanding our understanding of how chromatin effectors regulate DSB repair in plants.

## Introduction

To maintain genome integrity in the face of both endogenous and exogenous sources of DNA damage, organisms have evolved cellular programs to detect and repair different kinds of lesions ^1–3^. These programs are critical to prevent the accumulation of deleterious mutations that can reduce organismal fitness, resulting in developmental defects^2,4^ or the progression of diseases, including cancer^3^. These programs unfold within the context of chromatin—a biopolymer in which DNA is wrapped around histone proteins to form repeating arrays of nucleosomes^5^. For example, during double-strand break (DSB) repair, the DNA damage response (DDR) results in dynamic alterations of the chromatin surrounding the break as well as extensive transcriptional changes, both of which are critical for proper DNA repair^1,3,6–9^. While recent studies of plant responses to DSBs have shed light on the kinetics of gene expression changes in response to DNA damage^10^ and have demonstrated roles for several chromatin remodelling complexes^7,11^, our knowledge of how chromatin facilitates DNA repair in plants remains poorly understood, lagging far behind our understanding of these processes in yeast and animals.

While chromatin serves as the backdrop for DNA repair, it is not a uniform landscape. The positions, compositions, and compaction states of nucleosome arrays, as well as the histone proteins forming each nucleosome, can be modified by chromatin effectors to influence the accessibility and behaviour of the underlying genetic material^5^. These effectors can be broadly categorized into four groups: “Writers” that catalyze the addition of diverse chemical tags to specific residues within histone proteins, collectively termed histone modifications (*e.g.*, methylation, acetylation, phosphorylation, and ubiquitination)^12^. “Readers” that bind these modifications, “erasers” that catalyze their removal, and “remodelers” that use the energy of ATP to remove, slide, or exchange nucleosomes within chromatin^12^. In the context of DNA repair, some pathways are more reliant on chromatin effector complexes and chromatin modifications than others. For example, the repair of DSBs via canonical non-homologous end joining (NHEJ) pathways uses ligases to reseal unresected, broken DNA ends and thus, after the initial detection of DNA damage, this pathway requires relatively minimal alterations of chromatin^1,3,4^. On the other hand, chromatin modifications and chromatin remodeling complexes play important roles during the initial steps of repair via non-canonical NHEJ pathways and at many steps important for repair via homologous recombination (HR) pathways. In addition to facilitating the recognition and resection of DNA at breaks, specific chromatin modifications have been shown to mediate different repair steps, for example the recruitment of BRCA1 and RAD51 to promote homologous recombination^7,9,13–15^. While much less is known about roles of chromatin in these downstream repair steps in plants^7,15^, many aspects of the initial recognition of DSBs and signaling of the DDR are highly conserved.

Briefly, in plants, animals, and fungi, DSBs are recognized by conserved protein complexes that sense DNA damage and recruit serine/threonine kinases (*i.e.,* ATAXIA-TELANGIECTASIA MUTATED (ATM) and/or ATAXIA-TELANGIECTASIA MUTATED AND RAD3-RELATED (ATR))^1–3,7^. These kinases initiate signaling cascades through the phosphorylation of target proteins that regulate different aspects of the DNA damage response. These include, but are not limited to: phosphorylation of sensor proteins, which aid in the activation of ATM’s kinase activity^16^, phosphorylation of the histone variant H2A.X, which marks DNA repair foci and is one of the best characterized histone modifications involved in the DNA damage response^2,14^, and phosphorylation of important transcription factors (TFs), namely SUPPRESSOR OF GAMMA-RESPONSE 1 (SOG1) in *Arabidopsis*^2,17^ and p53 in mammals^18,1920^. Although not phylogenetically related, these master regulators of the DNA damage response share some target genes^10^ and serve similar functions in promoting genome stability by coordinating gene expression programs that promote DNA repair, modulate the cell cycle, and, in cases of extreme damage, initiate cell death^17,19,20^.

Another conserved feature of the DNA damage response is the involvement of complexes that remodel chromatin and alter histone acetylation levels at DSBs to facilitate DNA repair. In particular, the Nucleosomal Acetyltransferase of histone H4 (NuA4) and SWi2/snf2-Related (SWR1) complexes in yeast, as well as their SNF-2-related CREB-binding Activator Protein (SRCAP) and/or TIP60 counterparts in humans and drosophila, have interconnected and well characterized roles in regulating gene expression^21,22^ and coordinating DSB repair^23^. These multi-subunit complexes contain some distinct subunits, including the Essential SAS2-related Acetyltransferase 1 (Esa1) histone acetyltransferase in NuA4 and the SWi2/snf2-Related (Swr1) chromatin remodeler in SWR1^23^. However, they also have several shared accessory components, including Actin 1 (Act1), Actin Related Protein 4 (Arp4), Swr1 Complex 4 (Swc4), and Yeast ALL1-Fused gene from chromosome 9 (Yaf9)^23^. Together these unique and shared components facilitate the recruitment and activities of these remodelling complexes at chromatin. In the case of DSB repair in yeast, both the NuA4 and SWR1 complexes are recruited to break sites via the interaction of one of their shared subunits, ARP4, with ɣ-H2AX^23,24^. Once targeted, current models posit that the NuA4 complex acetylates H4 and H2A, as well as the histone variant H2AZ, which is incorporated by the remodelling activity of the SWR1 complex. These acetylation and histone exchange activities facilitate the generation of an open chromatin environment, promoting end resection and the recruitment of additional repair factors^23^. While the precise roles of many NuA4 and SWR1 components remain to be determined, mutants in key factors, including the catalytic subunits^25,2624,27,28^ and several of the shared proteins (*e.g.*, the YAF9^26–29^, SWC4^29^ and ARP4^24^ chromatin readers), are sensitive to various genotoxic stresses, demonstrating their important functions in the DNA damage response.

In plants, NuA4 and SWR1 complexes were identified using affinity purification approaches and shown to possess similar, though not identical, protein subunits as yeast, including orthologs of the catalytic subunits and all four of the shared proteins^30–37^. Interestingly, many of these subunits have been duplicated in plants^30^, suggesting they could act redundantly or serve specialized functions within their respective complexes. Like their yeast and animal counterparts, these plant complexes catalyse histone acetylation^30,38,39^ and H2AZ deposition^40–43^, respectively, and have demonstrated roles in regulating gene expression^30,42,43^. Specifically, they play important roles in controlling diverse biological processes including flowering time, reproduction, cell proliferation, and plant responses to the environment^30,42,43^. However, roles for these complexes in DNA repair are less well understood in plants. For the NuA4 complex, the *histone acetyltransferase of the myst family 1* (*ham1*) and *ham2* mutants are sensitive to UV-B damage^44^, but roles for these proteins and the NuA4 complex in DSB repair remain unexplored in plants. For the SWR1 complex, mutations in several SWR1-specific subunits, including the chromatin remodeler, *photoperiod-independent early flowering 1* (*pie1*), are sensitive to DSB-inducing agents and show defects in HR-like repair assays and in meiosis^45^. These findings demonstrate roles for chromatin remodelling complexes in plant DNA repair. However, many aspects of their functions in DSB repair remain to be explored, including how chromatin readers, like YAF9A and YAF9B, contribute to repair.

To identify chromatin effectors important for DNA repair in plants, a reverse genetic approach was utilized. While powerful, previous forward genetic screens for repair factors mainly identified Arabidopsis proteins important for NHEJ pathways, which are the preferred pathways utilized in somatic plant tissues^46^. This bias has resulted in a large gap in our understanding of how chromatin facilitates DNA repair via HR pathways in plants. To fill this gap, chromatin effectors co-expressed with genes important for DNA repair by HR, including BRCA1 and RAD51, were identified using a recently published transcriptional network of the Arabidopsis DNA damage response^10^. Among the co-expressed genes, YAF9B was identified. In Arabidopsis there are two YAF9 genes: *YAF9A*, which is constitutively expressed^32,47^, and *YAF9B*, which is induced by DNA damage^10^. Both YAF9A and YAF9B contain YEATS domains that bind acetylated histones^32^, confirming their roles as chromatin readers. While these proteins have redundant roles in regulating flowering time and plant development^31,32^, neither YAF9A nor YAF9B have known role(s) in plant DNA repair.

In the current work, we demonstrate that the induction of *YAF9B* expression in response to DNA damage is DSB specific and relies on a *cis*-regulatory promoter element bound by the SOG1 TF. We also show that both YAF9A and YAF9B are required for DSB repair using several different assays. In a “true-leaf” assay, which is most sensitive to defects in various NHEJ pathways, we found that YAF9A and YAF9B act redundantly to repair DSBs and promote meristem health (*i.e.*, its ability to produce new leaves). However, when we specifically assessed DNA repair via HR-like pathways using reporter lines, we found strong defects in *yaf9a* and *yaf9b* single mutants, revealing non-redundant roles. Based on these results, we hypothesize that YAF9A and YAF9B act redundantly in early steps of repair that are shared by many pathways (*i.e.*, end resection) but have distinct roles in later steps that are specific to HR-like pathways. To better understand how YAF9A and YAF9B function, we identified the protein complexes containing these chromatin readers. Consistent with previous reports^35^, YAF9A co-purified with components of both the NuA4 and SWR1 chromatin remodelling complexes. In contrast, YAF9B only co-purified with components of NuA4, defining a DNA damage-specific version of this complex. These findings not only provide the first evidence for a NuA4-specific component in mediating DSB repair in plants, but also shows that YAF9A and YAF9B are part of different complexes which, akin to the independent roles of the NuA4 and SWR1 complexes in DSB repair in yeast^23,48^, is in line with their non-redundant roles in HR-like repair. As both the plant NuA4 and SWR1 complexes have roles in transcriptional regulation under normal growth conditions, we assessed the effects of the *yaf9* mutants on the transcriptional response to DNA damage but found no defects. This supports our hypothesis that YAF9A and YAF9B play direct roles in DNA repair rather than indirect roles in regulating gene expression. Finally, we demonstrated that mutation of key residues within the reader domain of YAF9B are critical for its role in the DNA damage response. These results establish roles for YAF9A and YAF9B in DSB repair, highlighting the importance of histone readers and expanding our knowledge of how chromatin-associated factors govern DSB repair in plants.

## Results and Discussion

### *YAF9B* expression is DSB-specific, requires the SOG1 motif, and is the highest in meristematic tissues

The YAF9 family of YEATS domain chromatin readers^49–51^ were identified as candidates for regulating DSB repair in plants based on the expression pattern of the Arabidopsis *YAF9B* gene as well as roles for yeast YAF9 in promoting DNA repair and genome stability^52^. In Arabidopsis, the transcriptional response to treatment with DSB-inducing gamma-irradiation (γ-IR) was recently characterized over a 24 hour time course^10^. While *YAF9A* expression was high even under control conditions (- γ-IR) and was not altered by exposure to gamma-irradiation (+ γ-IR) (**Fig. 1A**), the expression pattern of *YAF9B* was similar to many genes important for DSB repair^10^: it showed low expression under control conditions but was strongly induced after exposure to γ-IR (**Fig. 1A**). Furthermore, SOG1, the master transcriptional regulator of the DNA damage response^10,53,54^, was enriched at its consensus motif within the *YAF9B* promoter (**Fig. 1A**)^10^, and *YAF9B* induction in response to DNA damage was lost in the *sog1-1* mutant (**Fig. 1A**)^10^. Together these observations suggest *YAF9B* is directly regulated by SOG1 during the DNA damage response.

**Figure 1:**
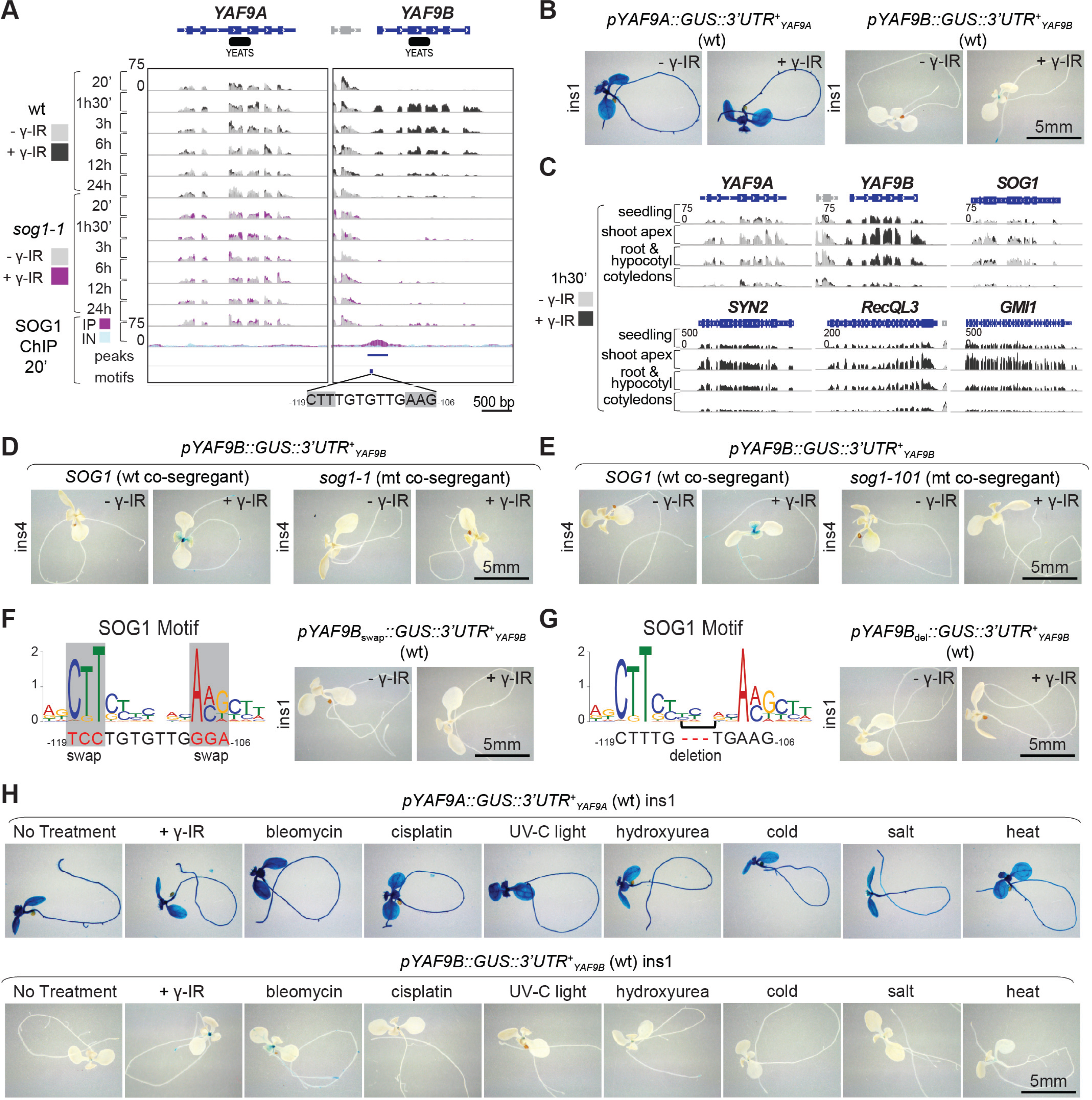
*YAF9B* expression is induced after DNA damage in a SOG1-dependent manner. (**A**) Genome browser tracks showing the expression levels of the YAF9A and YAF9B genes in 6-day-old wild-type (wt) or *sog1-1* mutant seedlings at different time points, specified in minutes (’) and/or hours (h), after mock (- γ-IR) or gamma-irradiation (+ γ-IR; 100Gy) treatments. SOG1 enrichment levels (+ γ-IR; 100Gy; 20’) from ChIP (IP) and input (IN) samples are shown below, including the SOG1 peak region in the YAF9B promoter, as well as the position and sequence (CTT-N_7_-AAG) of the SOG1 motif. The YAF9A and YAF9B gene models are blue, the neighboring genes are grey, and the position of the YEATS domains are black. The - γ-IR and + γ-IR mRNA-seq tracks and the IP and IN SOG1 ChIP-seq tracks are colored as indicated (left), superimposed, and shown on a scale of 0-75 reads per 10 million (RP10M). All data is from^10^. (**B, D, E, F, G**) Images showing the histochemical detection of β-glucuronidase (GUS) in 7-day-old seedlings 24h after mock (- γ-IR) or gamma-irradiation (+ γ-IR; 100Gy) treatments. For each group of images, the transgenes (insert (ins) numbers represent independent transgenic lines) and genetic backgrounds for the seedlings shown are as labeled. In panels **F** and **G**, the mutations introduced into the SOG1 motif in YAF9B promoter are indicated below the motif. (**C**) Genome browser tracks showing the expression levels of the indicated genes 1h30’ after mock (- γ-IR) or gamma-irradiation (+ γ-IR; 100Gy) treatments from either undissected 6-day-old seedlings or the indicated dissected tissues. The - γ-IR and + γ-IR mRNA-seq tracks are colored as indicated (left), superimposed, and shown in RP10M based on the indicated scales (upper left). (**H**) Images showing the histochemical detection of GUS in 7-day-old seedlings 24h after no treatment, exposure to gamma-irradiation (+ γ-IR; 100Gy), bleomycin (20 µg/mL), cisplatin (50µM), ultraviolet light (UV-C light,6000J/m2), hydroxyurea (80mM), salt (NaCl, 400mM), cold (4⁰C) or heat (37⁰C) treatments. For each group of images, the transgenes (including the insert number) and genetic backgrounds for the seedlings shown are as labeled.

To further support the connections between *YAF9B* expression, the DNA damage response, and the SOG1 transcription factor, *β-glucuronidase* (*GUS*) reporter constructs were generated. These reporters included either the endogenous *YAF9A* or *YAF9B* promoter and their respective *3’UTR*s flanking the *GUS* gene. Consistent with previous Reverse Transcriptase PCR (RT-PCR) data showing widespread expression of *YAF9A* under non-stressed conditions^32,47^, several independent *pYAF9A::GUS::3’UTR* reporter lines [insertions 1, 2 and 3 (ins1-3)] showed strong expression irrespective of γ-IR treatment across all tissues in both 7-day old seedlings (ins1; **Fig. 1B** and ins2- 3; **Fig. S1A**) and 14-day old seedlings (ins1-3; **Fig. S1A**). In contrast, the *pYAF9B::GUS::3’UTR* reporter lines showed expression only after γ-IR treatment, with the strongest *GUS* expression detected in the shoot apex and root tips of 7-day-old seedlings (ins1; **Fig. 1B** and ins2; **Fig. S1B**) and 14-day-old seedlings (ins1-2; **Fig. S1B**). Notably, the tissue specific *GUS* expression patterns observed for both the *pYAF9::GUS::3’UTR* reporter lines were consistent with the *in vivo* expression patterns for *YAF9A* and *YAF9B*, which were assessed directly via mRNA-seq experiments using dissected tissues from similarly staged wild-type (Col-0) seedlings either with or without exposure to γ-IR (**Fig. 1C**). While *YAF9A* was uniformly expressed across tissues and conditions, *YAF9B* expression was induced by γ-IR and was the highest in samples containing meristematic tissues (*i.e.* the shoot apex and root+ hypocotyl; **Fig. 1C**). These expression analyses confirm the broad, constitutive expression pattern of *YAF9A* and demonstrate that in addition to being induced by DNA damage, *YAF9B* expression is tissue-specific, mimicking both the expression pattern of the constitutively expressed *SOG1* gene^55^ and the damage responsiveness of several other DNA repair genes^56–59^ (**Fig. 1C**).

To assess the role of the SOG1 *cis*-regulatory motif in controlling the tissue-specific expression of *YAF9B,* the *pYAF9B::GUS::3’UTR* reporter was first crossed into two different *sog1* mutant backgrounds (*sog1-1*^54^ and *sog1-101*^59^) to confirm that the reporter recapitulates the SOG1- dependent expression pattern observed at the endogenous *YAF9B* locus (**Fig. 1A**). In both cases, the *pYAF9B::GUS::3’UTR* reporter was induced in meristematic tissues after γ-IR in the wild-type co-segregants and this expression was lost in the *sog1* mutants (**Fig. 1D, E**). Next, the role of the SOG1 *cis*-regulatory motif was assessed using mutated versions of the *pYAF9B::GUS::3’UTR* reporter where conserved bases within the SOG1 motif were swapped (purines for pyrimidines or vice versa) (**Fig. 1F**) or the spacing within the SOG1 motif was altered by deleting 3 bases (**Fig. 1G**). For each construct, several independent lines were tested, and in all cases mutations within the SOG1 motif resulted in strongly reduced *GUS* expression in both 7-day-old seedlings (ins1; **Fig. 1F, G** and ins2-3; **Fig. S1C, D**) and 14-day-old seedlings (ins1-3; **Fig. Fig. S1C, D**) as compared to the *pYAF9B::GUS::3’UTR* reporter with the unaltered SOG1 motif (**Fig. 1B and Fig. S1A**). These findings further support a direct regulation of *YAF9B* by SOG1 by demonstrating that the SOG1 *cis*-regulatory motif within the *YAF9B* promoters is required for *GUS* induction in meristematic tissues after DNA damage.

To assess the responsiveness of the *YAF9A* and *YAF9B* promoters to different types of DNA damage, the *pYAF9::GUS::3’UTR* reporter lines, along with wild-type lines as controls, were exposed to a variety of genotoxic conditions. For the *YAF9A* reporter, the GUS staining patterns were similar under all conditions (**Fig. 1H**), as was the expression of the endogenous YAF9A gene from wild-type plants (**Fig. S1E**), demonstrating that *YAF9A* expression is not significantly altered in response to any of these genotoxic stresses. For the *YAF9B* reporter lines, the GUS staining was specific to the shoot apex and root tips and was most prominent in response to DSB-inducing agents (γ-IR and bleomycin^57,60^) (**Fig. 1H**), with little to no expression observed in response to other stresses (*i.e.*, UV-C light for crosslinking^60–63^, cisplatin and hydroxyurea for replication stress^57,64^ and cold, salt and heat for environmental stress^57,65,66^) (**Fig. 1H**). Consistent with these results, expression of endogenous *YAF9B* was also the most strongly induced in response to DSB-inducing agents in wild-type plants (**Fig. S1E**). For all the aforementioned genotoxic stresses, the proper conditions were verified by assessing the induction of previously characterized stress-responsive genes^10,62,67–72^ (**Fig. S1F**). Taken together, these findings highlight the distinct expression patterns, *cis*-regulatory elements, and environmental responsiveness of the *YAF9A* and *YAF9B* genes.

### *yaf9* mutants are sensitive to genotoxic stress

To investigate the roles of YAF9A and YAF9B in DNA repair, several *yaf9* single and double mutants were characterized. Consistent with previous RT-PCR assays^32,47^, mRNA-seq experiments (with mock or γ-IR treatments) confirmed that the expression of *YAF9A* and *YAF9B* are clearly disrupted in the *yaf9a-1* and *yaf9b-2* mutants, respectively (**Fig. 2A**). In the *yaf9a-1* mutant, the transcript is truncated, and in the *yaf9b-2* mutant, the transcript is truncated and the expression is no longer responsive to γ-IR treatment (**Fig. 2A**). In contrast to these null mutants, the *yaf9b-3* mutant is weaker and condition specific, which is consistent with the position of the T-DNA in the *YAF9B* promoter between the SOG1 motif and the transcription start site (**Fig. 2A**). In a wild-type background *YAF9B* expression is in fact strongly reduced in the *yaf9b-3* mutant after γ-IR treatment (**Fig.2A**), suggesting that the proximity of the SOG1 motif to the *YAF9B* gene is important for *YAF9B* upregulation in response to DNA damage. However, the *yaf9a-1,b-3* double mutant (**Fig. S2A**) did not display the same developmental defects (*e.g.*, reduced plant size, infertility, etc.) previously observed in the *yaf9a-1,b-2* double mutant^31,32^ or another *yaf9a,b* mutant combination^73^, suggesting that the *yaf9b-3* mutant is weaker than the other *yaf9b* mutants. Indeed, in the *yaf9a-1,b-3* background, increased expression of the *yaf9b-3* allele was observed under both mock and γ-IR conditions (**Fig. 2A** and **Fig. S2B**). PCR amplification (**Fig. S2B**) and sequencing (**Fig. S2C** and **Source Data 1**) of the *YAF9B* transcript in the *yaf9b-3* single and *yaf9a-1,b-3* double mutants confirmed the presence of a full length, wild-type transcript, demonstrating that *yaf9b-3* is a knock down rather than a loss of function mutant, explaining its weaker developmental phenotype (**Fig. S2A**). Taken together, these *yaf9* alleles provide a series of mutants with variable strengths and expression patterns that can be leveraged to assess the roles of YAF9A and YAF9B in DNA repair.

**Figure 2:**
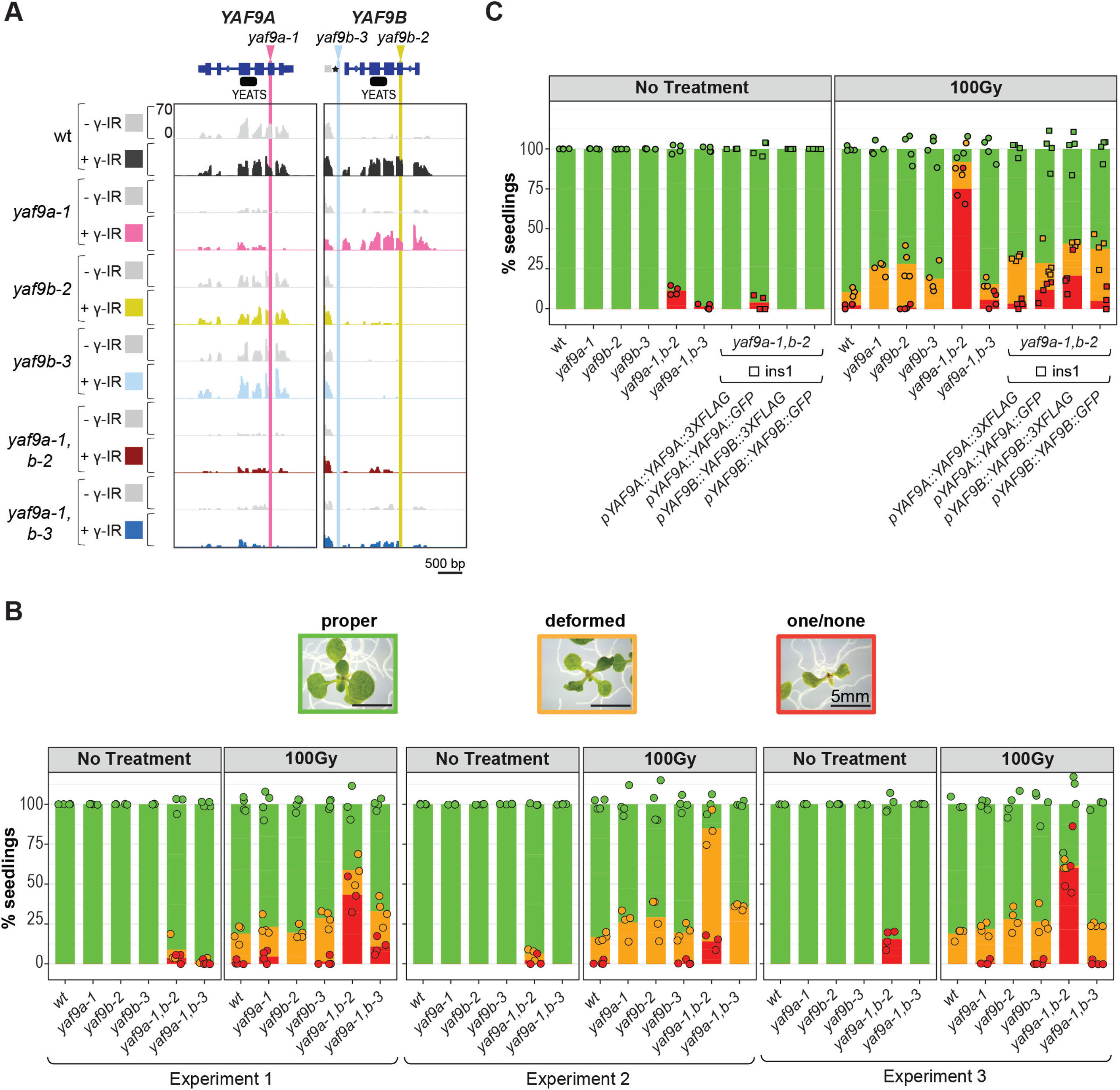
Genotoxic sensitivity in *yaf9* mutants. (**A**) Genome browser tracks showing the expression levels of YAF9A and YAF9B in 8-day-old seedlings 3 hours after mock (- γ-IR) or γ- irradiation (+ γ-IR; 100Gy) treatments in the genotypes indicated on the left. The - γ-IR and + γ- IR tracks for each genotype are colored as indicated and shown on a scale of 0-70 RP10M. The gene models for YAF9A and YAF9B are marked with the T-DNA positions of the yaf9a-1 (pink), yaf9b-2 (mustard), and yaf9b-3 (blue) alleles and the location of the YEATS domains are shown in black. The location of the SOG1 motif in the YAF9B promoter is indicated with an asterisk (*). (**B** and **C**) True leaf assays showing the percentage of 11-day-old seedlings categorized as having proper true leaves (green), deformed true leaves (orange) or one/none true leaves (red) after either no treatment or treatment with 100Gy. Images of representative seedlings for each category are shown in above the bar plots in panel **B**. For each bar plot, the genotypes are labeled on the x-axis. On the y-axis, the color-coded dots represent the percentage of seedlings belonging to each category calculated based on sets of ∼30 seedlings that represent biological replicates (n=3 or 4). The stacked bars are the average of all the data points for each category. In **C**, the data from one insert (ins1) of each tagged construct are shown as squares, with data for additional inserts (ins2-3) shown in **Fig. S2F** and **G**.

Using the aforementioned *yaf9* mutants, defects in DNA repair were first assessed using the “true leaf” assay. In this assay, the accumulation of DNA damage is quantified by counting the number of deformed or missing true leaves that arise in young seedlings exposed to genotoxic stresses due to cell cycle arrest and/or cell death in the apical meristem^45,74^. Compared to both wild-type and no treatment controls, a higher proportion of the *yaf9a-1,b-2* double mutant plants showed true leaf defects after exposure to 100 Gy of γ-IR across three independent experiments (**Fig. 2B**) and these defects arose in a dose-dependent manner (**Fig. S2D**). The other single and double *yaf9* mutants behaved similar to the wild-type control (**Fig. 2B**), demonstrating redundancy between YAF9A and YAF9B and revealing that even the low level of *YAF9B* expression in the *yaf9a-1,b-3* double is sufficient to promote plant tolerance to genotoxic stress. To confirm that the damage sensitivity observed in the *yaf9a-1,b-2* mutant is due to disruption of the *YAF9A* and *YAF9B* genes, complementation assays were conducted using transgenes containing the *YAF9A* or *YAF9B* genes driven by their endogenous promoters and tagged with either 3XFLAG- or GFP. To avoid the fertility defects observed in the homozygous *yaf9a-1,b-2* double mutant, these transgenes were introduced into either *yaf9a-1(ho),b-2(het)* or *yaf9a-1(het),b-2(ho)* lines, respectively, as diagrammed in **Fig. S2E**. After selecting for homozygous *YAF9A* and *YAF9B* transgenic lines, and genotyping for the *yaf9a-1* and *yaf9b-2* alleles, true leaf assays were conducted. For these assays, six independent YAF9A and YAF9B tagged lines (three with 3XFLAG (ins1-3) and three with GFP (ins1-3)) were utilized (ins1; **Fig. 2C** and ins2-3; **Fig. S2F, G**). As is evidenced by the reductions in seedlings with deformed or missing true leaves after γ-IR exposure, these assays demonstrated that both the 3XFLAG- and GFP-tagged YAF9A and YAF9B proteins were able to largely rescue the DNA damage defects observed in the *yaf9a-1,b-2* mutant (**Fig. 2C** and **Fig. S2F, G**), confirming that YAF9A and YAF9B act redundantly to promote plant tolerance to genotoxic stress.

### YAF9A and YAF9B have non-redundant roles in DSB repair via homologous recombination

Given the co-expression of *YAF9B* with genes required for homologous recombination (HR)^10^, two previously described HR-like reporters^75,76^ were utilized to directly assess the roles of YAF9A and YAF9B in DSB repair. Both these reporters (**Fig. 3A**) contain a *GUS* gene that is nonfunctional until a DSB is generated at a specific location (red tick mark) and then repaired either by single strand annealing (SSA) at the DGU.US reporter or by synthesis-dependent strand annealing (SDSA) at the IU.GUS reporter. To achieve high rates of HR-like repair at these reporters, an inducer construct that expresses the I-SceI restriction enzyme and generates DSBs in the *GUS* reporter at the desired location is required (**Fig. 3A**). However, lower rates of repair are also observed after exposure to random mutagenesis, including γ-IR exposure (**Fig. 3A**). In all scenarios, restoration of the *GUS* gene through HR-like repair allows rates of SSA and SDSA to be determined by counting the number of blue sectors in plants harboring these reporters.

**Figure 3:**
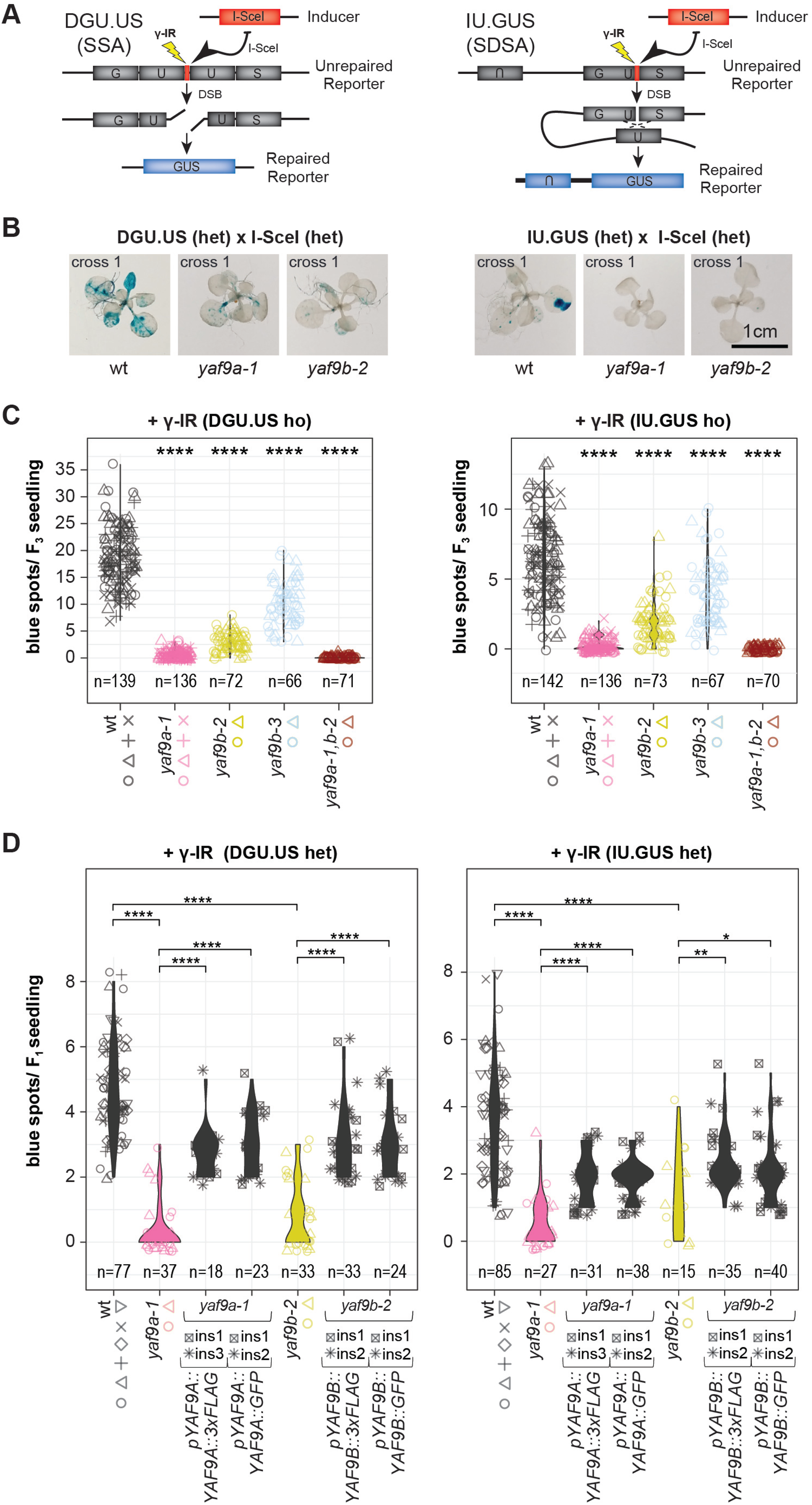
YAF9A and YAF9B are required for homologous recombination after DSB damage. (**A**) Transgenic DSB reporter constructs (DGU.US/SSA and IU.GUS/SDSA)^75,76^ in which a β-glucuronidase (GUS) gene is interrupted by a unique sequence including the 18-bp I-SceI recognition site (red bar). DSBs can be introduced into theses GUS reporters via 2 methods: They can be induced precisely at the unique I-SceI sequence by introduction of the I-SceI enzyme on a separate transgene or they can be introduced at a lower rate by exposure to genotoxic agents, like γ-IR (lightning bolt), that introduce DSBs randomly throughout the genome. In either case, repair of the DSB via HR-like mechanisms generates a functional GUS gene that can be detected as blue spots after histochemical staining. (**B**) Representative images showing the recombination events via SSA/DGU.US and SDSA/IU.GUS in 15-day-old F_1_ seedlings homozygous for the genotypes indicated below each image and heterozygous for the reporter and inducer constructs indicated above. Detailed crossing schemes are shown in **Fig. S3A**. Multiple crosses were performed and the cross number is indicated on the top-left corner of the image, with images from additional crosses shown in **Fig. S3B, C**. (**C** and **D**) Violin plots showing the recombination events via SSA/DGU.US and SDSA/IU.GUS after 100Gy γ-IR in seedlings with the genotypes indicated below and the reporters indicated above each graph. For the DGU.US and IU.GUS reporters recombination events were scored in 14-day-old or 19-day-old seedlings, respectively. In **C**, the data points for F_3_ seedlings from different lineages (i.e., different F_2_ parents of the indicated genotypes; see **Fig. S3A** for crossing details) are represented by the different color-coded shapes. In **D**, the data points for F_1_ seedlings from different lineages (i.e., the progeny of different genetic crosses; see **Fig. S3D** for crossing details) are represented by the different color-coded shapes. For each epitope-tagged YAF9 constructs two independent inserts (ins) are shown. For **C** and **D**, the asterisks (*) represent the p-value from Wilcoxon tests comparing each genotype to the wt control (not significant (ns): p > 0.05; *: p <= 0.05; **: p <= 0.01; ***: p <= 0.001; ****: p <= 0.0001).

To test the effects of *yaf9* mutants on DSB repair using the aforementioned reporters, the required genetic materials were generated via the crossing schemes outlined in **Fig. S3A**. Using the inducer system, the number of blue sectors resulting from repair of either the DGU.US/SSA or IU.GUS/SDSA reporters were visibly lower in the *yaf9a-1* and *yaf9b-2* seedlings when compared to seedlings from the wild-type control crosses (**Fig. 3B**). Furthermore, this phenotype was consistent across several independent crosses (**Fig. S3B, C**). To assess the effects in additional mutant backgrounds, these assays were also conducted in the absence of the inducer, with damage generated by exposure to γ-IR (see **Fig. S3A** for crossing scheme). In these assays, the number of blue spots resulting from HR-like repair were significantly reduced in all the *yaf9* mutants compared to the wild-type controls in both reporter backgrounds (Wilcox t-test **** p ≤ 0.0001; **Fig. 3C**). In line with *yaf9b-3* being a knockdown rather than a null allele, this mutant displayed a weaker phenotype than *yaf9b-2* (**Fig. 3C**). By comparison, the recombination rates in the *yaf9a-1* mutant were lower than both *yaf9b* single mutants (**Fig. 3C**), revealing a stronger repair defect in this mutant. However, the repair defects in the *yaf9a-1,b-2* double mutant were only slightly enhanced compared to the *yaf9a-1* single mutant (**Fig. 3C**). These findings demonstrate that, unlike for the true leaf assay (**Fig. 2B, C**), YAF9A and YAF9B cannot fully compensate for each other and instead act in a largely non-redundant manner to promote HR-like repair of DSBs.

To confirm that the DSB repair defects observed in the *yaf9* mutants are due to reduced recombination rates, rather than, for example, epigenetic silencing of the *GUS* reporter lines^76,77^, complementation assays were conducted. For these assays, the DGU.US/SSA and IU.GUS/SDSA reporter lines, in either the wild-type, *yaf9a-1*, or *yaf9b-2* backgrounds used in **Fig. 3C**, were crossed with the previously generated YAF9 complementing lines (**Fig. S2E**), and recombination rates were assessed in the resulting F_1_ seedlings after exposure to γ-IR (**Fig. 3D**, see **Fig. S3D** for crossing scheme). Compared to the uncomplemented *yaf9a-1* and *yaf9b-2* controls, recombination rates were significantly increased in the presence of multiple, independent tagged YAF9 proteins (**Fig. 3D**), demonstrating a partial rescue of the recombination defects at both HR-like reporters. This complementation affirms that the *GUS* reporters were not epigenetically silenced and remain competent for recombination and GUS detection in these *yaf9* mutant backgrounds. Taken together, these findings rule out indirect effects of YAF9A and YAF9B in transgene silencing, affirming their critical roles in DSB repair via HR-like mechanisms.

### YAF9A and YAF9B facilitate DSB repair in a chromatin-mediated but transcription-independent manner

The DNA damage response involves the regulation of thousands of genes, many of which are critical for DSB repair itself or for other repair-associated biological processes like cell cycle regulation (**Fig. 4A**)^10,66,67,78,79^. To determine whether the roles for YAF9A and/or YAF9B in DNA repair are due to changes in gene regulation upon exposure to DNA damage, three replicate mRNA-seq experiments [batch (b) 1-3] were performed using wt or *yaf9* mutant seedlings 3 hours after either mock (- γ-IR) or γ-irradiation treatments (+ γ-IR; 100Gy) (**Table S1**). As an initial control, differentially expressed genes (DEG; |Fold change (FC)| ≥2 and p-value ≤0.01) were identified using just the wt seedling data +/- γ-IR treatments ([wt (+ γ-IR)]_b1-3_ vs. [wt (- γ-IR)]_b1-3_; **Fig. 4B** and **Table S2**). These analyses recapitulated the previously published transcriptional changes^10^, with similar patterns for the up- and downregulated genes at the 3 hour time point (*i.e.*, path 1 genes are the most upregulated in response to γ-IR, followed by path 2, *etc*.). Furthermore, these control samples were robustly separated by condition (+/- γ-IR) in a principal component analysis (PCA) across all three experiments (b1-3; **Fig. 4C**). In this PCA, a clear separation between the +/- γ-IR samples was also observed for all the *yaf9* single and double mutants (**Fig. 4C**), suggesting the DNA damage response remains largely intact. Furthermore, with the exception of the *yaf9a-1,b-2* mutant, which displays strong developmental defects even under normal conditions^31,32^, all the other *yaf9* mutants clustered together with their respective wild-type controls (**Fig. 4C**).Together, these analyses demonstrate wild-type-like expression profiles under both mock and DNA damage conditions for these *yaf9* mutants.

**Figure 4:**
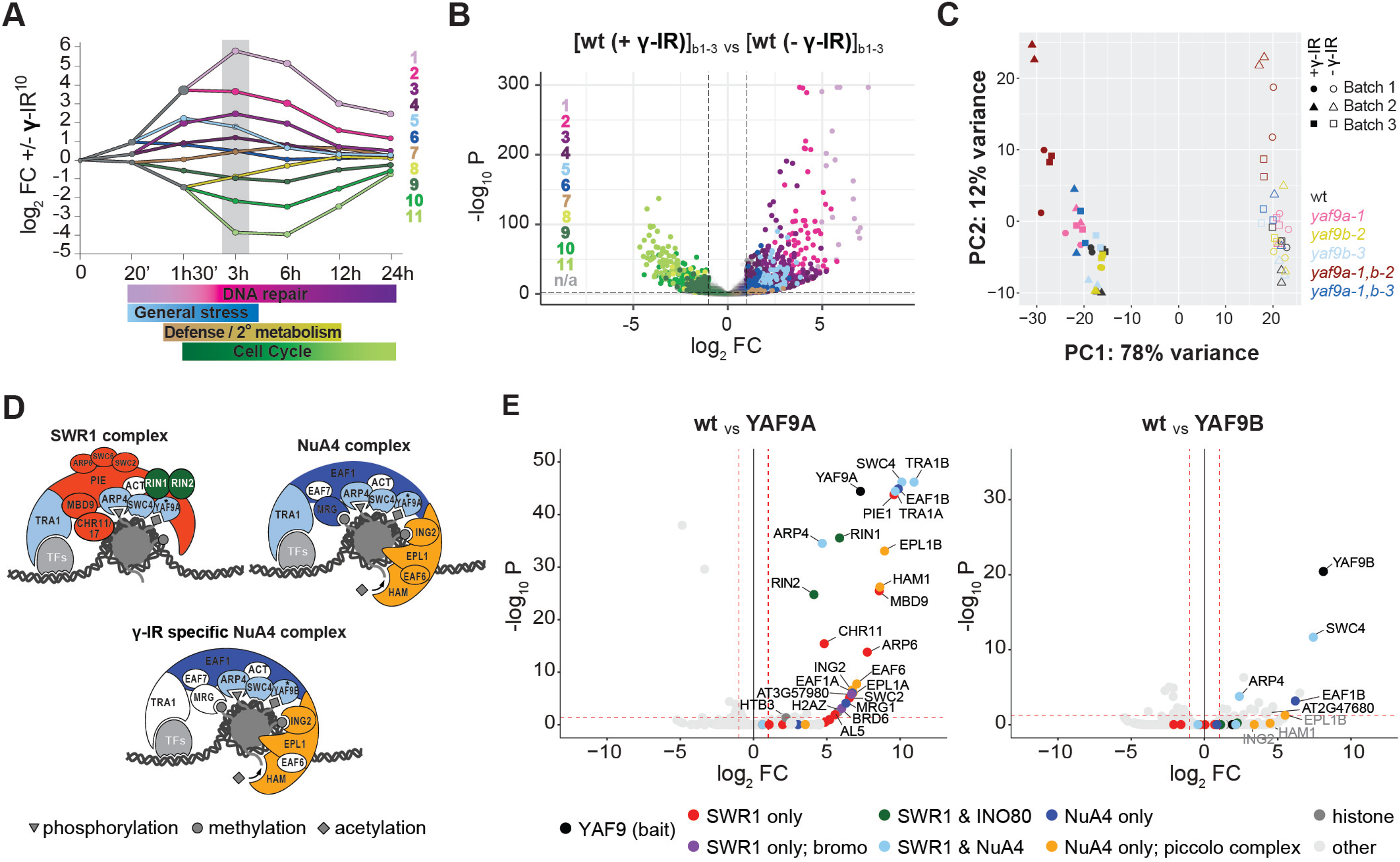
YAF9A and YAF9B are present in distinct chromatin remodeling complexes and facilitate DNA repair via mechanisms that are independent from roles in transcriptional regulation. (**A**) Reproduction of the DREM model from^10^, summarizing the transcriptional response of Arabidopsis seedlings exposed to DNA damage. The x-axis shows the log_2_ FC in expression of genes after exposure to γ-irradiation (+ γ-IR; 100Gy) or mock (- γ-IR) treatments. The genes are grouped together into 11 differently colored paths (1-11) based on their expression patterns across a time course (y-axis) where the time is shown in minutes (’) and hours (h). The number (n) of genes in each path are indicated and the major gene ontology terms for select paths (indicated by their corresponding colors) are shown below. The 3h time point, which was selected for analysis of the roles for the YAF9 proteins in the DNA damage response, is highlighted in grey. (**B**) Volcano plot showing the changes in gene expression observed in wild-type seedlings 3 hours after exposure to DNA damage. The x- and y-axes show the -log_10_ P and log_2_ FC values for DEGs identified by comparing the genotypes and conditions indicated above the plot across three independent mRNA-seq experiments [batches (b)1-3]. DEGs present in the DREM model shown in **A** are color coded based on the gene groups in which they reside. DEGs not present in the DREM model are shown in gray. The dotted lines represent |log_2_ FC| ≥1 and p-value ≤0.01. (**C**) Principal component analysis (PCA) showing data points from all 3 batches mRNA-seq experiments. The different experiments (batch 1-3) are shown in different shapes, the different genotypes are shown in different colors, and the different conditions (3h after 100Gy; + γ-IR or 3h after mock; - γ-IR) are shown as filled or open shapes, respectively. For each genotype and condition, two technical replicates were included. (**D**) Cartoons based on Bieluszewski *et al*.^111^, summarizing the current understanding of the *Arabidopsis* SWR1^35–37^ and NuA4^33^ complexes, color-coded and labeled with sub-components. Filled subunits were enriched via YAF9A (upper) or YAF9B (lower) IP-MS experiments. (**E**) Volcano plots showing the enrichment or depletion of proteins from a combination of several independent YAF9A or YAF9B IP-MS experiments compared to non-transgenic, wild-type IP-MS experiments. The red hashed lines demarcate a |log2 FC| ≥1 and an p-value ≤0.05. Enriched proteins previously identified as components of the SWR1 and NuA4 remodeling complexes are labeled and colored based on the legend below. Proteins labeled in grey are enriched but feel below the adjusted p-value cut-off.

The global observations from the PCA were also confirmed via DEG analyses using the *yaf9* mutants (|FC| ≥2 and p-value≤0.01; **Table S2**): Under mock conditions [*mutant* (- γ-IR) vs. wt (- γ-IR); **Fig. S4A**], very few genes were mis-regulated in the *yaf9* single or *yaf9a-1,b-3* double mutants, none of which play known roles in DNA repair (**Table S2**). Consistent with the PCA analysis, many more genes were mis-regulated in the *yaf9a-1,b-2* double mutant (**Fig. S4A;** 61 down, 513 up DEGs and **Table S2**). However, after DNA damage [*mutant* (+ γ-IR) vs. *mutant* (-γ-IR); **Fig. S4B**], all the *yaf9* single and double mutants showed a transcriptional response similar to the wild-type control (**Fig. S4B** vs. **Fig. 4B**). In fact, these responses were so similar that comparisons between the mutant and wild-type γ-irradiation datasets [*mutant* (+ γ-IR) vs. wt (+ γ- IR); **Fig. S4C**] revealed very few mis-expressed genes. Among all the *yaf9* mutants, only the *yaf9a-1,b-2* double showed a significant difference in its DNA damage response (**Fig. S4C;** 132 down, 1131 up DEG), but this difference corresponds to a slightly more, rather than less, robust response when compared to the wild-type control. This is evident by the larger scale of expression changes observed for DNA damage response genes in the *yaf9a-1,b-2* mutant in both the volcano plot shown in **Fig. S4C** and in the heatmaps shown in **Fig. S4D** [*i.e.*, some genes normally upregulated after DNA damage (paths 1-7) are more upregulated in the *yaf9a-1,b-2* mutant, and some genes normally downregulated (paths 8-11) are more downregulated in the *yaf9a-1,b-2* mutant, but the converse situation (genes normally up, that are now down *etc.*) is not observed]. Taken together, these analyses demonstrate that the transcriptional response to DNA damage is not negatively impacted in either the *yaf9* single or double mutants. Furthermore, since the *yaf9a* and *yaf9b* single mutants have strong defects in DSB repair (**Fig. 3**) but show little to no changes in gene expression under normal conditions or after exposure to DNA damage (**Fig. S4**), these data support direct roles for YAF9A and YAF9B in DNA repair, akin to roles proposed for yeast YAF9^52^, rather than indirect roles in regulating the transcriptional response to DNA damage.

To determine whether YAF9B, like YAF9A^35^, interacts with NuA4 and SWR1 chromatin-modifying complexes (**Fig. 4D**), which could facilitate their roles in DNA repair^45^, immunoprecipitation and mass spectrometry (IP-MS) experiments were performed. These experiments utilized 8-day-old seedlings from non-transgenic wild-type controls and representative 3XFLAG-tagged YAF9A or YAF9B complementation lines, in their respective single mutant backgrounds (*pYAF9A:YAF9A:3XFLAG/yaf9a-1* (ins1) and *pYAF9B:YAF9B:3XFLAG/yaf9b-2* (ins1); **Fig. S2E and 3D**). For YAF9A one IP-MS experiment was conducted under mock conditions and another 6 hours after exposure to 100Gy of γ-IR. For YAF9B, two IP-MS experiments were also conducted, both 6 hours after exposure to 100Gy of γ- IR. As the replicate experiments for YAF9A and YAF9B gave similar results (**Table S3**), they were analyzed together to identify enriched copurifying with either YAF9A or YAF9B proteins, respectively (**Fig. 4D, E**). Consistent with previous reports, YAF9A copurified with nearly all the previously identified NuA4 and SWR1 components (**Fig. 4D, E,** and **Table S3**). However, the YAF9B purification suggests a preferential, if not specific, interaction with the NuA4 complex. When considering the most significant co-purifying factors (|log_2_ FC >1| and adjusted p-value <= 0.05), as well as those that are highly enriched (log_2_ FC > 3) despite less significant adjusted p-values, YAF9B only interacts with shared NuA4/SWR1 proteins and NuA4-specific proteins, but no SWR1-specific proteins (**Fig. 4D, E** and **Table S3**). These include SWC4 and ARP4 from the shared targeting module, but notably not YAF9A. Indeed, even in the YAF9A experiment conducted after exposure to γ-IR, YAF9B was not detected (**Table S3**). These findings suggest that each complex only contains a single YAF9 subunit, which is consistent with previous results showing both YAF9A and YAF9B interact with the same subunit, SWC4, by Y2H^31^. In addition to these shared components, YAF9B also co-purified with another protein previously determined by Y2H, EAF1B^31^. This NuA4-specific protein serves as a scaffold for the shared NuA4/SWR1 targeting module as well as the NuA4-specific piccolo catalytic complex, for which EPL1B, HAM1, and ING2 were all enriched in the YAF9B purification. In contrast, the SWR1-specific proteins showed little to no enrichment. This interaction data demonstrates that YAF9B forms a distinct, DNA damage-specific NuA4 complex. Taken together with the absence of transcriptional defects in *yaf9a* and *yaf9b* mutants (**Fig. 4C** and **Fig. S4**), the roles for YAF9A and YAF9B in DNA repair (**Figs. 2, S2, 3,** and **S3**), and past roles demonstrated for other SWR1-complex components in DSB repair^45^, these findings link YAF9A and YAF9B, and their respective chromatin remodeling complexes, to DSB repair in a manner that is chromatin-mediated but transcription-independent.

### Conserved residues within the YEATS domain are required for YAF9B function

To investigate whether the acetylation reader function of the YAF9B YEATS domain^32^ is important in mediating its role in DNA repair, a mutagenesis approach was employed. The Arabidopsis YAF9A and YAF9B proteins share homology with human GAS41 and yeast Yaf9^30,52^. The residues required for the YEATS domain to recognize lysine acetylation in GAS41 have been identified and shown to form an aromatic cage^80,81^ (**Fig. 5A**; green triangles). Based on an alignment with GAS41 (**Fig. 5A**) and a structural prediction of YAF9B^82,83^ (**Fig. 5B**), amino acids H89, S91, F92, W111, G112 and F114 were identifed as candidate residues important for the YAF9B YEATS domain-histone interaction (**Fig. 5A-C**). To assess the roles of these amino acids in DNA repair, alanine substitutions of the aforementioned residues, either alone or in combinations of up to four residues, were introduced into both the 3XFLAG- and GFP-tagged YAF9B constructs and transformed into *yaf9a-1(het),b-2(ho)* plants. Proper protein expression and accumulation of the wild-type or mutated 3XFLAG-tagged YAF9B proteins were verified using western blots (**Fig. S5A, B**). True leaf assays were then performed in *yaf9a-1,b-*2 double mutants homozygous for either the wild-type or the mutated YAF9B transgenes (**Fig. 5D, E**). Consistent with previous assays (**Fig. 2B**), the sensitivity in *yaf9* single mutants after γ-IR was comparable to the wild-type control, the *yaf9a-1,b-2* double mutant was more sensitive, and addition of functional YAF9B rescued the true leaf defects (**Fig. 5D, E**). Amongst the single amino acid substitutions, the S91A, F92A, G112A and F114A single mutants all complemented the *yaf9a-1,b-2* double mutant to a similar level as observed for the wild-type version of YAF9B (**Fig. 5D, E**), demonstrating that alone, these mutations are not sufficient to disrupt YAF9B function *in vivo*. In contrast, the YAF9B constructs harboring the W111A mutation all showed lower levels of complementation, with the defects in the W111A,G112A,F114A triple and the H89,W111A,G112A,F114A quadruple mutant combinations nearly equivalent to the *yaf9a-1,b-2* double mutant (**Fig. 5D, E**). This evidence suggests that these residues play an essential role in the YEATS domain function. Notably, the sensitivities observed in the YAF9B substitutions are in agreement with the previously identified importance of their counterparts in GAS41^84^. Moreover, the importance of the Tryptophan (W) in the aromatic cage for YEATS domain function in both *Arabidopsis* YAF9B and Human GAS41^84^, suggests a conservation of the histone-binding mechanism. Altogether, this data demonstrates that a functional YEATS domain is required for the role of YAF9B in DNA repair.

**Figure 5:**
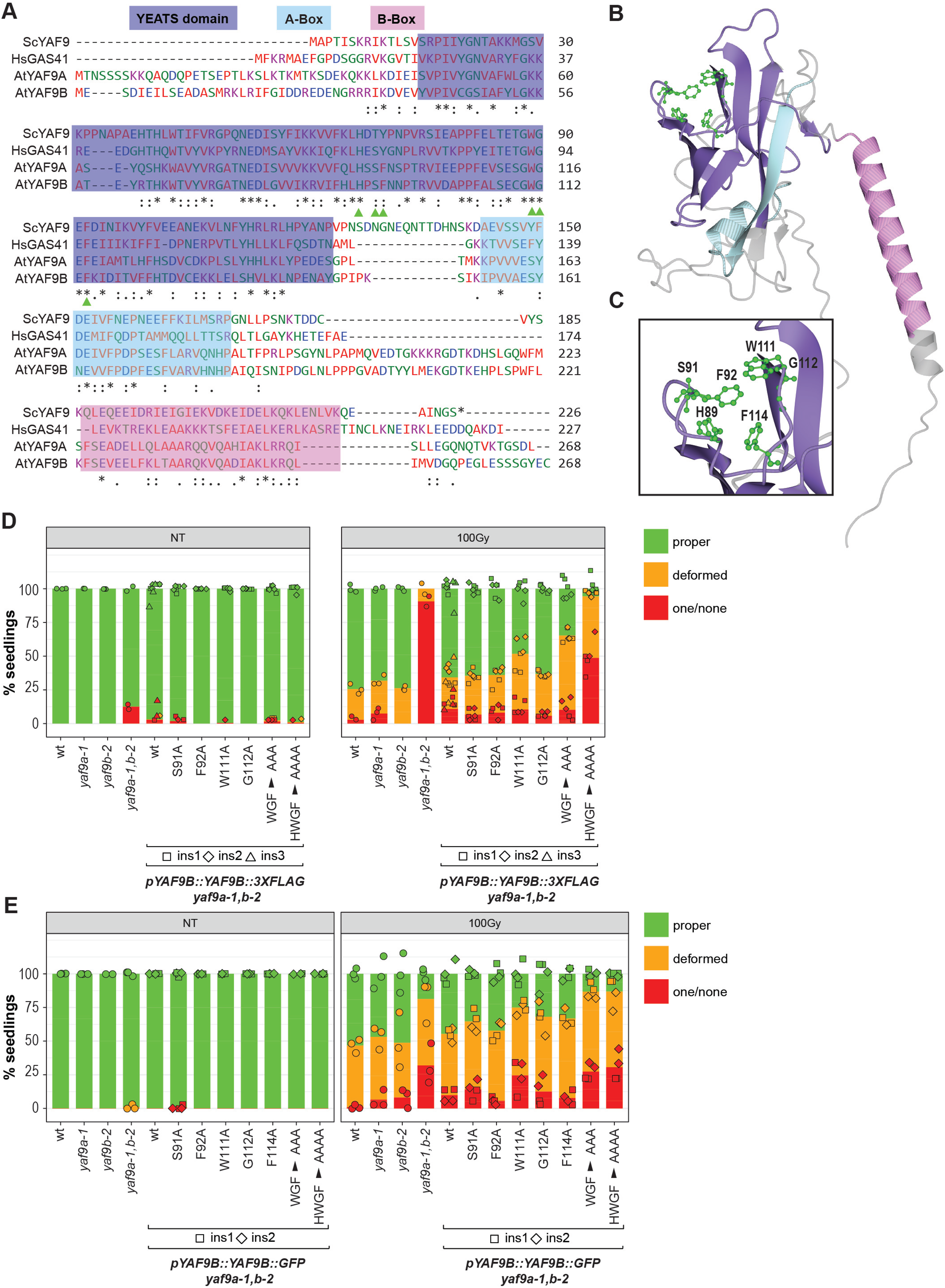
The YEATS reader domain is required for YAF9B function. CLUSTALW alignment^112^ of ScYAF9, HsGAS41, AtYAF9A and AtYAF9B. The YEATS domain is highlighted in dark blue, the A-box in light blue, the B-box in pink. The residues required for the YEATS domain to recognize lysine acetylation in GAS41^113^ are indicated with green arrows and the “.”, “:”, and “*” symbols denote weakly conserved residues, strongly conserved residues and fully conserved residues, respectively. (**B**) AlphaFold predicted 3D-structure of YAF9B^82,83^ where the YEATS domain, A-Box, and B-Box are colored as in (**A**). The residues predicted to interact with histone acetyl marks are shown in green (ball and stick) are labeled with the amino acid symbol and number, with a zoomed view depicted in (**C**). True leaf assays showing the rescue capacity of YAF9B with key acyl mark interactors mutated using (**C**) 3XFLAG- and (**D**) GFP-tagged lines. The plot shows the percentage of seedlings with proper true leaves (green), deformed true leaves (mustard) and one/none true leaves (red) in 12-day-old seedlings after treatment with 100Gy in yaf9 single, double mutants and rescue lines. Each rescue line is represented by 2-3 biological replicates (independent insertions, represented by different point shapes) and 2-3 technical replicates (seeds from same seed packet grown on different plates and randomized positions in the growth chamber). Each color-coded dot represents the percentage of seedlings belonging to one of the three true leaf phenotype categories calculated based on ∼30 seedlings. Each stacked bar is the mean of 3-4 data points represented by dots.

## Conclusion

In this work, we expand the roles of YAF9A and YAF9B to include not just plant development and reproduction^31,32,47^ but also DSB repair. First, we determined the DNA damage- and tissue-specific regulation of *YAF9B* by the SOG1 transcription factor via a *cis* element present in the *YAF9B* promoter (**Fig. 1** and **S1**). Second, we demonstrated roles for YAF9A and YAF9B in promoting plant meristem health in response to genotoxic stress (**Fig. 2** and **S2**) and facilitating DSB repair via HR-like mechanisms (**Fig. 3** and **S3**). Third, we found that YAF9A and YAF9B denote distinct chromatin remodeling complexes (**Fig. 4D, E** and **Table S3**), with YAF9B defining a DNA damage-specific NuA4 complex while YAF9A, like its yeast counterpart, associates with both the SWR1 and NuA4 complexes^23,30^. This finding represents the first link between the NuA4 complex and DSB repair in plants and suggests a plant-specific innovation to cope with DNA damage—the formation of a specialized NuA4 complex that is enriched in meristematic tissues. Finally, we connected the function of YAF9B in DNA repair to its YEATS reader domain (**Fig. 5** and **S5**). Together, these findings expand our understanding of the chromatin effectors that regulate DSB repair in plants and reveal both conserved and plant-specific aspects of YAF9 functions.

Mechanistically, our studies revealed that the *yaf9a* and *yaf9b* mutants have no effect on the transcriptional response to DNA damage (**Fig. 4** and **S4**) but show defects in several different DNA repair assays (**Fig. 2, 3** and **S2, 3**), supporting direct roles in DNA repair rather than indirect roles through gene regulation. While the precise roles of YAF9A and YAF9B during DSB repair require further investigation, their associations with the SWR1 and NuA4 complexes (**Fig. 4D, E** and **Table S3**), and the behaviors of the *yaf9a* and *yaf9b* mutants in different types of repair assays, offer clues to their functions. For example, in the true leaf assay, which scores many different types of repair (*e.g.*, both NHEJ and HR pathways), YAF9A and YAF9B act in a largely redundant manner as only the *yaf9a,b* double mutants show strong defects upon exposure to DNA damage (**Fig. 2** and **S2**). However, in the GUS reporter assays, which specifically assess repair via HR-like pathways^85^, YAF9A and YAF9B are largely non-redundant as both the *yaf9a* and *yaf9b* single mutants show reduced rates of repair (**Fig. 3** and **S3**). While these results are seemingly contradictory, our understanding of the SWR1 and NuA4 chromatin remodeling complexes associated with these YAF9 proteins suggests a plausible explanation.

The activities of the SWR1 and NuA4 complexes are interconnected and associated with the promotion of open chromatin states in yeast, mammals, and plants^23,30,42^. The NuA4 complex catalyzes the acetylation of lysine residues in the N-terminal tails of H4, H2A, and H2A.Z^23,30^, which neutralizes their positive charge and thus weakens their interactions with the negatively charged DNA backbone^86^. Likewise, the SWR1 complex facilitates the incorporation of the H2A.Z variant^23,42,43^, which confers lower nucleosome stability in some conditions ^87^. Within the context of DNA repair, the formation of open chromatin promotes end resection, an early DNA processing step that is important for non-canonical NHEJ and HR pathways^9,85,88^. Based on these functions, we hypothesize that the roles of *Arabidopsis* YAF9A and YAF9B are redundant for early DNA repair steps, like end resection, that are shared between non-canonical NHEJ and HR pathways. For the latter steps in repair, the yeast NuA4 and SWR1 complexes (or the combined mammalian TIP60 complex) also play important roles, as the acetylation of histone tails and the incorporation of the H2A.Z variant facilitate the recruitment of additional factors to DSBs to promote HR-based repair^88–90^. In plants, we hypothesize that YAF9A and YAF9B, and their associated SWR1 and NuA4 complexes, play non-redundant roles in these later steps of HR-based repair since loss of either YAF9A or YAF9B results in strong repair defects (**Fig. 3** and **S3**). Furthermore, as both YAF9A and YAF9B associate with NuA4 complexes (**Fig. 4D, E** and **Table S3**), yet the *yaf9b* single mutant shows strong defects in DNA repair (**Fig. 3** and **S3**), we posit that the YAF9A-NuA4 and YAF9B-NuA4 complexes also have distinct roles in DNA repair. To further support these hypotheses, it will be important to determine what specific roles each of these YAF9 proteins/complexes play in early and late stages of DSB repair and how they regulate chromatin to promote HR-like mechanisms.

As a first step towards understanding the functions of the Arabidopsis YAF9 proteins at chromatin, we investigated the importance of the YAF9B YEATS domain for DNA repair. YEATS domains are histone binding domains initially characterized as readers of lysine acetylation. However, more recent work in yeast and mammals has shown YEATS domains can also bind, and even prefer, other types of lysine acylations, including propionylation, butyrylation, and crotonylation, that fit within the same binding cleft^50,51^. In Arabidopsis, both YAF9A and YAF9B were found to bind acetylated histone tails^32^. However, their preferences for binding acetylation at different lysine positions or within different histone tails (H3, H4, H2AZ) remains limited and their potential to bind other types of acylations remains completely unknown. Despite many questions remaining about the specificity of the Arabidopsis YAF9 YEATS domains, mutation of highly conserved aromatic residues predicted to form π-stacking interactions with acyl-lysines residues revealed that the YAF9B YEATS domain is critical for its role in DNA repair. This finding marks a major advance in our understanding of how chromatin modifications coordinate DNA repair in plants and represents one of the few direct links between DNA repair, in this case DSB repair, and a chromatin reader function (*i.e.*, the YAF9B acyl-lysine activity). Although the importance of their reader domains in repair has not been explicitly tested, other notable examples of such connections include PDS5C and MSH6, which have Tudor binding specificities^83,91^ and repair defects^92,93^ that strongly support direct roles in mismatch and homology-dependent repair at genes marked with H3K4me1, respectively. Likewise, BCP4 and XIP, which have BRCA1 C-terminal (BRCT) domains shown to bind γ-H2A.X^94,95^, play important roles in DSB repair as revealed by true leaf and HR-like assays, respectively^94,95^. This growing list of connections between chromatin readers and DNA repair pathways/steps represent initial, but critical advances towards understanding how chromatin modifications coordinate repair to ensure genome stability.

In the case of YAF9B, the importance of its YEATS domain for DNA repair suggests a role either in the targeting or retention of the NuA4 complex at chromatin in response to DNA damage. To fulfill this role, several non-mutually exclusive scenarios can be envisioned. For example, the YEATS domain of YAF9B could be required to recruit the NuA4 complex to chromatin surrounding DSBs with pre-existing acylated histones. Alternatively, NuA4 recruitment could instead depend on ARP4-mediated recognition of phosphorylated H2AX (a mark that rapidly accumulates at DSBs via the kinase activities of ATM and ATR^2,14^), as previously demonstrated in yeast^24^. Once recruited the NuA4 complex could then acetylate the surrounding chromatin, generating binding sites for the YAF9B YEATS domain and facilitating the stable accumulation and/or spreading of NuA4 at DSBs. Or, a combination of these two mechanisms could be required to facilitate NuA4 targeting and efficient DNA repair. Likewise, the YAF9A YEATS domain could be required for the recruitment and/or retention of the SWR1 complex at DSBs to facilitate the incorporation of H2A.Z and the subsequent repair of DSBs. In moving forward, gaining a better understanding of the specificities of the YAF9 YEATS domains and the order of action for the different YAF9 complexes will be especially important in distinguishing between the proposed scenarios and revealing how the YAF9 chromatin readers coordinate different steps of DSB repair.

## Materials and methods

### Plant Materials

All *Arabidopsis* mutant and transgenic lines used in this study were in the Col-0 ecotype and were grown at 22°C under long-day conditions (16h light, 8h dark) unless specified otherwise. The following previously characterized transfer DNA (T-DNA) insertion mutants were obtained from *Arabidopsis* Biological Resource Center (ABRC): *yaf9a-1* (SALK_106430^47^), *yaf9b-2* (SALK_046223^31^), and *sog1-101* (GABI_602B10^59^). The *sog1-1* EMS mutant^54^, which is in a mixed Landsberg erecta (Ler)/Col background containing a *pCYCB1;1::GUS* fusion, was provided by A. Britt, Department of Plant Biology, University of California, Davis, CA. A second, uncharacterized *yaf9b* allele, *yaf9b-3* (WISCDSLOX377-380P19/CS853514^96^), was obtained from ABRC. Homozygous lines were identified by PCR-based genotyping using the primers listed in **Table S4**.

### Developmental phenotyping

Developmental phenotypes of plants grown in the greenhouse (22°C under long-day conditions: 16h light, 8h dark) shown in **Fig. S2A** and **E** were captured using a digital camera. Images were processed using photopea (https://www.photopea.com/).

### Generation of gateway plasmids and transgenic *Arabidopsis* plant lines

Constructs containing *GUS* reporters or epitope-tagged versions of *YAF9A* or *YAF9B* were generated using the MultiSite Gateway® Three- Fragment Vector Construction Kit (Cat # 12537- 023, Invitrogen). To generate BP constructs containing either the promoter regions, genic regions (without STOP codon), or 3’UTR regions of *YAF9B* and *YAF9A*, the corresponding genomic regions were amplified using Phusion® High-Fidelity DNA Polymerase (Cat # M0530, NEB) from wild-type genomic DNA using the primers listed in **Table S4**. The resulting PCR products were gel purified (Cat # D4001, Zymo Research) and recombined into the following pDONR entry vectors (pDONR-P4P1r for promoter regions, pDONR-221 for genic regions, and pDONRP2rP3 for 3’-UTR regions as detailed in **Table S5**) using the Gateway BP Clonase II Enzyme kit (Cat # 11789020, Invitrogen) according to manufacturer’s instructions. These sequence-verified BP clones, as well as previously generated BP clones containing (pDONR221-GUS, pDONRP2rP3- 3XFLAG-BLRP^mut^ and pDONRP2rP3-GFP) were then recombined with the pB7m34GW destination binary vector^97^ using the Gateway LR Clonase kit (Cat# 11791019, Invitrogen) to generate the constructs detailed in **Table S5**.

Constructs containing mutations in the SOG1 motif or in the YEATS domains of YAF9A or YAF9B were generated by performing Site Directed Mutagenesis (SDM) on the BP constructs pDONRP4-P1R-pYAF9B, pDONR221-YAF9A, or pDONR221-YAF9B, respectively, using the primers listed in **Table S4**. SDM was performed using a two-step method. Briefly, the original BP constructs were amplified with Phusion® High-Fidelity DNA Polymerase (Cat # M0530, NEB) and complimentary primers containing the desired mutation (listed in **Table S4**). The PCR product was digested with DpnI (Cat # R0176S, NEB) and transformed into One Shot™ MAX Efficiency™ DH5α-T1R Competent Cells (Cat # 12297016, Invitrogen). These BP clones were sequence verified for the SDM mutations and were used to generate the destination vector constructs listed in **Table S5**.

After sequence verification, the destination vectors were transformed into the AGLO *Agrobacterium tumefaciens* strain, and used to create *Arabidopsis* transgenic lines in specified genetic backgrounds (**Table S5**) using the floral dip method^98^. Primary transformants for each construct were selected on half strength Linsmaier and Skoog (LS½) medium with 25ug/mL Basta (Cat # J66186, Alfa Aesar) and genotyped for the genetic background. Subsequent generations were screened to select homozygous, single insert lines in the appropriate genetic backgrounds.

Lines containing the *pYAF9B::GUS:: 3’UTR^+^*construct in *sog1* mutant backgrounds were generated by crossing the *pYAF9B::GUS:: 3’UTR^+^*line (ins4, which was generated in a *yaf9a-1 (het) yaf9b-2 (ho)* background), into either the *sog1-1* or *sog1-101* mutant. In the F_3_ generation, lines homozygous for the *pYAF9B::GUS::3’UTR^+^*reporter, wild-type for *yaf9a-1* and *yaf9b-2*, and then either mutant or wildtype for the *sog1* allele were identified and utilized as co-segregation pairs for GUS assays. For crosses with the *sog1-1* mutant a *pCYCB1;1::GUS* transgene present in this mutant line was also segregated out by genotyping.

### GUS histochemical staining of seedlings

Seeds were sterilized in microcentrifuge tubes by exposure to chlorine gas (200mL bleach and 5mL hydrochloric acid (HCl), stirred continuously) in a sealed container for 1h and then aired out for 1h to vent the chlorine gas. Sterilized seeds were resuspended in 1mL of sterilized water, stratified by incubation at 4°C for 2-3 days in the dark, sowed on LS½ media plates and grown in long day conditions (16h light, 8h dark). Whole seedlings (8-day-old and 14-day-old for promoter driven *GUS* lines in **Fig. 1, S1** and 14-day and 19-day old for HR-like repair assays in **Fig. 3, S3**) were fixed by submersion in ice-cold 90% acetone for 20 minutes. The seedlings were then briefly rinsed by submersion in GUS staining buffer (50mM NaPO4, pH7.2, 0.2% Triton X-100) at room temperature and then incubated in GUS staining buffer supplemented with 2 mM X-Gluc (Cat # G1281C1, GoldBio) at 37°C in the dark overnight (10-16h). The seedlings were cleared by sequential ethanol washes (30%, 50% and then 70% ethanol) for 1h each. The cleared seedlings were stored in 70% ethanol and imaged using a stereo microscope (FisherbrandTM Research Grade Stereo Zoom Microscope) and photographed with a SeBaCam14C digital camera (SEBACAM14C CMOS 5V 14mp CMOS, Laxco) using SeBaView digital imaging software (Laxco). Larger seedlings were imaged on a lightbox and photographed with a digital camera. The recombination efficiency for **Fig. 3C, D** was quantified by counting the number of blue spots in the shoot of the seedling using a stereo microscope (FisherbrandTM Research Grade Stereo Zoom Microscope) at 1.5X magnification.

### RNA expression experiments RNA isolation

For all three *yaf9* mRNA-seq experiments (see **Table S1**), seeds were chlorine gas sterilized, stratified at 4°C for 3 days in the dark and grown on LS½ media with 0.6% plant agar and 1% sucrose for 8-days in long day conditions (16h light/ 8h dark cycles at 22°C). The seedlings were then either mock treated or exposed to γ-IR (100Gy) at a dose rate of 8-10Gy/min using a Co60 radioactive source and then returned to long day conditions for 3h. Two replicates of 6-8 whole seedlings from the mock and γ-IR treated samples were collected, frozen in liquid nitrogen, and stored at -80°C. Total RNA was extracted using the Quick-RNA MiniPrep kit (Zymo Research R1055) according to the manufacturer’s protocol.

For the tissue dissection mRNA-seq, 6-day-old seedlings, grown vertically in constant light conditions (24h light at 22°C) were used. Approximately 100 mock or γ-IR (100Gy) treated seedlings were collected 1h30’ post treatment and fixed in chilled (-20°C) 80% acetone and infiltrated under vacuum for 5’ twice. The seedlings were dissected individually using a dissection scope (VWR VistaVision Stereo Microscope) in 100% ethanol with fine forceps and collected in 2mL microcentrifuge tubes filled with 100% acetone. For the RNA extraction, the acetone was completely aspirated prior to tissue lysis using liquid nitrogen and a Tissue Lyser (Qiagen), and total RNA was isolated using the RNeasy Micro Kit (Qiagen 74004) according to the manufacturer’s protocol.

### mRNA-seq Experiments

#### Library preparation and sequencing

For all mRNA-seq experiments, mRNAs were purified from 1-2µg of total RNA using the NEBNext Poly(A) mRNA Magnetic Isolation Module (New England Biolabs E7490), mRNA libraries were prepared with the NEBNext UltraII RNA Library Prep Kit (New England Biolabs E7770) and sequenced on Illumina HiSeq 2500 (50bp single-end mode).

#### Data processing

The time course mRNA-seq data for the wild-type and *sog1-1* after mock or γ- IR (100Gy) treatments and the SOG1-3xFLAG ChIP-seq data were previously published^10^ and available in Gene Expression Omnibus (GEO) database (GSE112773). The mRNA-seq data from the current study is available in Gene Expression Omnibus (GEO) database. For all mRNA-seq experiments, Illumina reads were mapped to the TAIR10 genome using STAR^99^, with the following parameters: maximum number of mismatches per read = 2, minimum intron length = 20 bp, maximum intron length = 6000 bp, minimum total length of exons = 5% of read length. A summary of the read mapping is presented in **Table S1**. Downstream analyses were conducted using the HOMER suite as follows^100^. The mapped SAM files were converted to BAM using SAMtools^101^ and were used to generate Tag directories using the makeTagDirectory script from HOMER^100^. To visualize the gene expression data, UCSC browser tracks and .tdf files were generated for all mRNA-seq samples using the makeUCSCfile script (-fragLength given -norm 10000000 -style rnaseq -strand both) from HOMER^100^ and igvtools^102^, respectively.

Expression values for all genes across samples were retrieved using the analyzeRepeat.pl script from HOMER^100^ with the -noadj and -condenseGenes options (**Table S2A**). Using this table, PCA plots were generated (with expression values listed in **Table S2A**) in R-Studio using plotPCA function of DESeq2^103^ package using experimental design parameters: design = ∼batch + Condition + Group + Condition:Group. The *yaf9* mRNAseq data was corrected for batch effects using removeBatchEffect function from limma package^104^. Differentially expressed genes (FC ≥ 2 with an FDR ≤ 0.01) based on the experimental conditions (i.e., + γ-IR vs. – γ-IR or mutant vs. wild-type) were identified using the a script from HOMER^100^ with the -DESeq2^103^ and -repeats options. The list of differentially expressed genes in each comparison are listed in **Table S2B** and the log_2_FC and adjusted p-value for all genes are listed in **Table S2C**. The Volcano plots in **Fig. 4B** and **S4A-C** were generated using the EnhancedVolcano package^105^. Cut-offs of log_2_FC >|1| and p-value < 10e^-2^ were used and genes were colored based on the previously identified DDR gene group (based on DREM model) they belonged to ^10^. The heatmaps in **Fig. S4D** were generated using the pheatmap package^106^ based on the log_2_FC values generated during the DESeq2 analysis. Individual heatmaps were generated for each group of genes present in the DREM model.

### Gel-based expression analysis of the *YAF9B* transcript

1.5µg of total RNA (YAF9 mRNA-seq, set4) was utilized for cDNA synthesis using the High-Capacity cDNA Reverse Transcription Kit (Cat#4374967, Life Technologies) according to manufacturer’s instructions. The cDNA was used as a template and RT-PCR was performed with GoTaq® Green Master Mix (M7123) and primers listed in **Table S4** in a thermocycler (95°C for 3min, followed by 32 cycles of denaturation at 95°C for 30s, annealing at 57°C for 30s and elongation at 72°C for 1min, followed by 72°C for 5min and 12°C forever). The resultant PCR product was migrated on a 1.5% agarose Tris-Acetate-EDTA (TAE) gel alongside a DNA ladder and imaged on a Biorad Gel Doc.

### Stress treatments

For all stress treatments, seeds (Col-0 and the YAF9 promoter driven GUS lines) were chlorine gas sterilized, stratified, sown onto a nylon mesh on LS½ media plates with 8g/L plant agar and grown vertically for 7 days in in long-day conditions (16h-light/ 8h-dark cycles). After 7 days, the plates were split into mock and treatment sets. For the treatment set, the nylon meshes and seedlings were treated as follows: For the γ-IR and UV-C light stress, the seedlings were moved to fresh LS½ media plates with 8g/L plant agar and then exposed to 100Gy γ-IR (dose rate of 8- 10Gy/minute using a Co60 radioactive source) or 6000J/m2 UV-C light (dose rate of approximately 5000J/m2/minute using a Strategene UV stratalinker). Treated seedlings were moved back to long-day growth chamber. After 3h of recovery 4-6 Col-0 seedlings were collected for RNA extraction and expression analysis and after ∼24h of recovery 5-15 *pYAF9::GUS::3’UTR* seedlings were collected for GUS staining as previously described. For the heat and cold treatments, the seedlings were moved to fresh LS½ media plates with 8g/L plant agar and the plates were placed vertically either in 37°C chamber (dark) or 4°C cold room for 3h. Immediately following the 3h treatment, 4-6 Col-0 seedlings were collected for RNA extraction and expression analysis and after the 24h treatment, 5-15 *pYAF9::GUS::3’UTR* seedlings were collected for GUS staining as previously described. For bleomycin, cisplatin, hydroxyurea and salt treatments, the seedlings were moved to freshly made LS½ media plates with 8g/L plant agar supplemented with 20µg/mL bleomycin (SIGMA 15361, stock= 2mg/mL in DMSO), 50µM cisplatin (SIGMA P4394, stock = 1.67mM in water), 80mM hydroxyurea (SIGMA H8627, stock = 30mg/mL in water) or 400mM NaCl (stock = 5M NaCl in water) respectively. These plates were moved back to long-day growth chamber. After 3h of treatment on these drug supplemented plates 4-6 Col-0 seedlings were collected for RNA extraction and expression analysis and after ∼24h of treatment 5-15 *pYAF9::GUS::3’UTR* seedlings were collected for GUS staining as previously described. For the mock treatment set, the seedlings were moved to LS½ media plates with 8g/L agar as in all cases and grown in the long-day chamber, with all tissue collections in parallel to the corresponding treatment set for each stress.

For relative gene expression analysis, total RNA was isolated post treatment (as detailed above) from 4-6 Col-0 whole seedlings, ground to a powder in liquid nitrogen, using the Quick-RNA MiniPrep kit (Cat #R1055, Zymo Research) according to manufacturer’s interactions. RT-qPCR was performed on cDNA (synthesized as described above) with Bio-Rad CFX384 Real-Time System using Luna Universal qPCR Master Mix (Cat# M3003E, NEB) according to manufacturer’s instructions. Transcript levels were determined using the standard curve method and relative expression was calculated by normalizing to a control gene, *AT5G13440*. The experiments were performed several times, and each experiment included 2 technical replicates. The gene-specific primers used in this analysis were listed in **Table S4**.

### True leaf assay

*Arabidopsis* seeds were surface sterilized using the bleach-HCl method and stratified in sterile water at 4°C in dark for 3 days. The stratified seeds were γ-IR treated at a dose rate of 8-10 Gy/min using a Co60 radioactive source, sown onto LS½ plates with 1% sucrose and 0.6% agar, and then transferred to a growth chamber with 16h light/ 8h dark cycles at 23°C. After 11-12 days, the plants were scored if they have proper true leaves, deformed true leaves or one/no true leaves. The percentage of seedlings in each category was visualized using a stacked bar chart generated using ggplot2 package^107^ in R-Studio. The example true leaf phenotypes for seedlings with proper true leaves, deformed true leaves and one/no true leaves (shown in **Fig. 2B)** were imaged using stereo microscope (FisherbrandTM Research Grade Stereo Zoom Microscope) and photographed with SeBaCam14C digital camera (SEBACAM14C CMOS 5V 14mp CMOS, Laxco) using SeBaView digital imaging software (Laxco).

### Analysis of recombination via SSA and SDSA

The genetic materials used for assessing recombination at the SSA and SDSA reporters were generated as diagramed in **Fig. S3A** and **Fig. S3D** and recombination rates were assessed by GUS staining (See GUS histochemical staining of seedlings). For assays utilizing the I-SceI inducer construct to generate DSBs (**Fig. 3B, S3B, C**), GUS staining was performed on F_1_ seedlings grown for 15-days in long-day conditions (16h light, 8h dark). GUS-stained seedlings were imaged on a lightbox using a digital camera and processed using photopea (https://www.photopea.com/). For assays using γ-IR to generate DSBs, 7- or 8-day-old seedlings (**Fig. 3C, D**; DGU.US and IU.GUS, respectively) were grown in long-day conditions (16h light, 8h dark), exposed to 100Gy γ-IR, and let recover under long-day conditions until GUS staining was performed on 14- or 19-day-old seedlings (**Fig. 3C** DGU.US and IU.GUS or **Fig. 3D** IU.GUS, respectively). The number of blue spots per seedlings were counted using a stereo microscope (FisherbrandTM Research Grade Stereo Zoom Microscope) using 1.5X magnification. P-values were calculated from Wilcox t-tests between the indicated genotypes on R-Studio (compare_means function in package ggpubr^108^).

### Western Blotting

For western blots (**Fig. S5**), four 8-day-old seedlings were ground to a fine powder in liquid nitrogen 3 hours after mock or 100Gy γ-IR treatments. The powder was resuspended in 100 µL 2X LaemmLi and boiled for 15 minutes. Samples were migrated on a Criterion Bis-Tris 10% gel (BioRad #3450112), transferred in a CAPS buffer (10 mM CAPS, 10% v/v methanol, pH adjusted to 11 with NaOH) to a 0.45 PVDF blotting membrane (Cat # 10600023, Amersham), blocked overnight at 4°C with 5% w/v milk in TBS-T (Tris-Buffered Saline with 0.1% Tween 20), incubated with HRP-conjugated anti-FLAG antibodies (1:10,000; SIGMA A8592) in 1% w/v milk in TBS-T (1h at room temperature), washed in TBS-T (three times for 10’ each) and detected using the SuperSignal West Dura Extended Duration Substrate (Cat # 34075, Thermo Scientific) according to manufacturer’s instructions. Ponceau S staining of Rubisco (∼50 kDa) serves as a loading control. For Ponceau-S staining, PVDF membranes were incubated in Ponceau-S solution (5% acetic acid (v/v), 0.2% Ponceau-S (w/v)) for 5 minutes and were de-stained using deionized water.

### Immunoprecipitation and Mass Spectrometry

For both YAF9A YAF9B replicates, *Arabidopsis* seeds (*pYAF9B::YAF9B::3XFLAG yaf9b-2*, *pYAF9A::YAF9A::3XFLAG yaf9a-1*, and negative control Col-0) were sterilized first in a 70% ethanol solution containing 0.05% w/v SDS for 10 minutes and then in a 90% ethanol solution for 5 minutes. The sterilized seeds were sprinkled on LS½ media plates with 8g/L agar, stratified at 4°C for 2-3 days in the dark and grown vertically in a growth chamber with 16h light/ 8h dark cycles at 23C for 8 days. For both YAF9B replicates, 8-day-old seedlings from each genotype were exposed to 100Gy γ-IR (8-10Gy/min using a Co60 radioactive source), moved back to long-day growth chamber till sample collection (6h after treatment), and then frozen in liquid nitrogen. For YAF9A and its negative control, one replicate was exposured to 100Gy γ-IR using the same conditions as for the YAF9B experiments, while the other replicate was conducted without exposure to γ-IR. For the immunoprecipitation assays, 10g of frozen tissue was ground in liquid nitrogen and resuspended in 50mL of lysis buffer (LB: 50mM Tris pH7.6, 150mM NaCl, 5mM MgCl2, 10% glycerol, 0.1% NP-40, 0.5mM DTT, 1µg/µL pepstatin, 1mM PMSF and 1 protease inhibitor cocktail tablet (Roche, 14696200)). The tissue was then homogenized by douncing and centrifuged at 4°C in an ultracentrifuge for 25 minutes at 13,500 rpm. Each supernatant was incubated at 4°C for 1hwith 250μL of anti-FLAG magnetic beads (Cat # M8823, SIGMA) by gentle rotation on a HulaMixer. The FLAG beads were then washed twice with 20mL of LB and five times with 1mL of LB. For each wash, the beads were rotated at 4°C for 5 minutes. Proteins were then released from the FLAG beads during five room temperature incubations with 400µL of 3XFLAG peptide (Cat # F4799, SIGMA) at a concentration of 100µg/mL in Phosphate buffered saline (PBS). The eluted proteins were precipitated using trichloro acetic acid (TCA). Samples were digested with trypsin and analyzed using a Thermo Fusion Lumos mass spectrometer couped to a Dionex Ultimate 3000 UHPLC using a data-dependent acquisition strategy as described in Felgines *et al.*^109^. The MaxQuant output files were subsequently processed for statistical analysis of differentially enriched proteins using the IPinquiry4^110^ R package and visualized as volcano plots using the ggplot2^107^ R package. The raw data for both batches of IP-MS data are available through the MassIVE repository via the identifier MSV000099158.

## Acknowledgements

We thank colleagues and lab members for their comments on the manuscript and Dr. Holger Puchta for the HR reporter lines. J.A.L. was supported by the Rita Allen Foundation and a gift from the Hess Corporation. N.V. was supported by Salk Women in Science Award, Jesse and Caryl Philips Foundation Fellowship and Chapman Charitable Trust Graduate Student Scholar Award. C.B. was supported by the LL Nippert Charitable Foundation and Catharina Foundation. AM.S.P. was supported by a postdoctoral fellowship from the Paul F. Glenn Center for Biology of Aging Research at the Salk Institute. This work was also supported by the NGS Core Facility of the Salk Institute with funding from NIH-NCI CCSG: P30 CA01495, NIH-NIA San Diego Nathan Shock Center P30 AG068635, the Chapman Foundation and the Helmsley Charitable Trust. The Illumina sequencing datasets generated in this study were deposited in the NCBI Gene Expression Omnibus (GEO) and are accessible via the accession number GSE308666.

---

**Figure S1:**
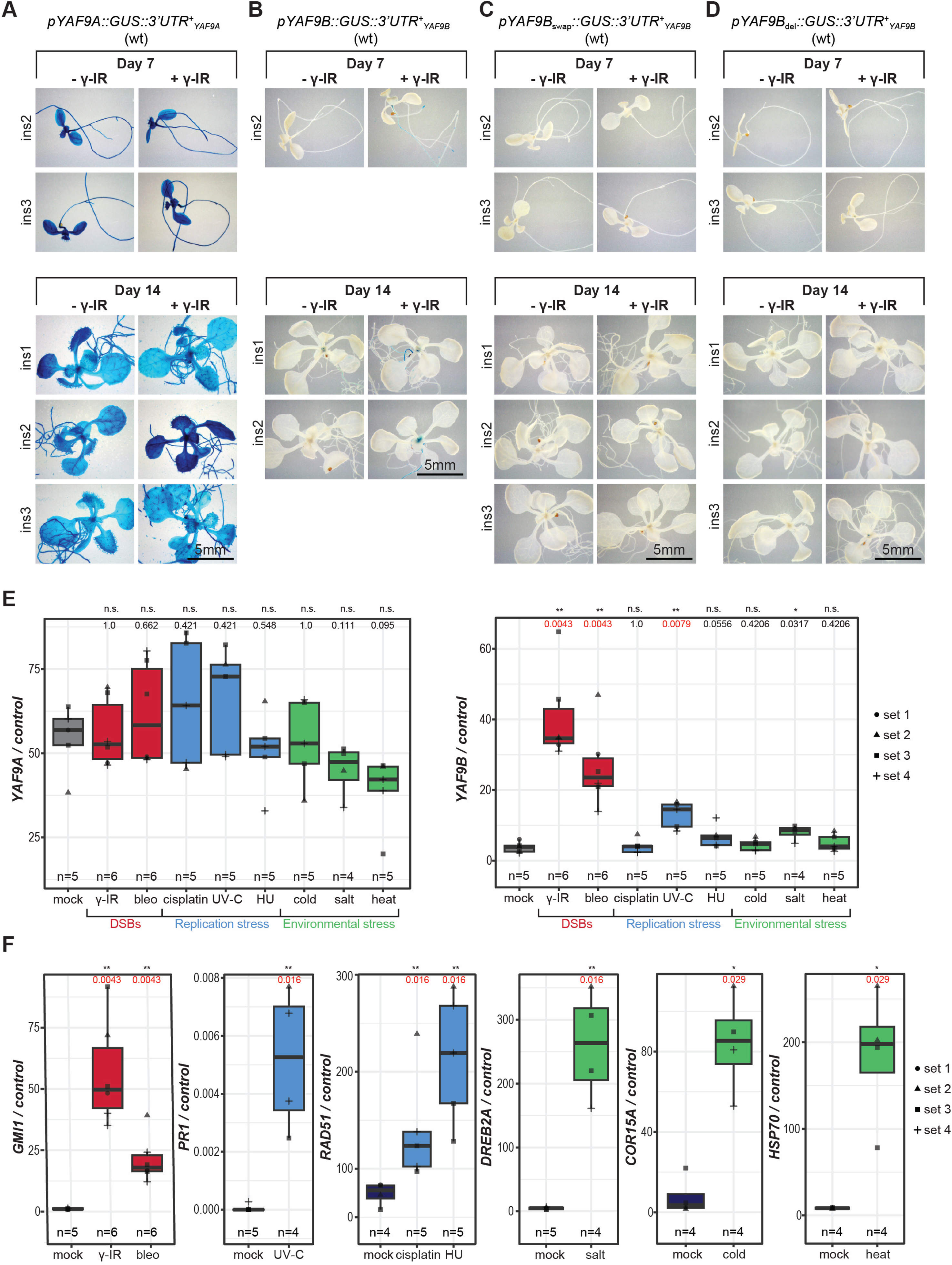
*YAF9B* expression is induced after DNA damage in a SOG1-dependent manner. (**A-D**) Images showing the histochemical detection of GUS in 7- and 14-day-old seedlings 24h after mock (- γ-IR) or gamma-irradiation (+ γ-IR; 100Gy) treatments. For each group of images, the transgenes (including the insert number) and genetic backgrounds for the seedlings shown areas labeled. (**E** and **F**) Boxplots from several independent RT-qPCR experiments (sets 1-4) assessing the relative expression levels of YAF9B and YAF9A (**E**) or genes known to be induced by different types of DNA damage (**F**). Specifically, GMI1 for γ-IR^10^ and bleomycin^67^; PR1 for UV-C^68^; RAD51 for cisplatin^62^ and hydroxyurea^69^; DREB2A for salt stress^70^; COR15A for cold stress^71^; and HSFA6A for heat stress^72^. For all experiments, expression levels were normalized to a control gene, AT5G13440, and samples were collected from 7-day-old wt seedlings 3h after exposure to the stress treatments indicated on the x-axis. Each data point represents the average of two technical replicates and the shapes correspond to the four independent experiments (sets 1-4). The total number (n) of data points included for each condition is indicated and conditions in which the expression levels of the YAF9 genes (**E**) or various stress responsive genes (**F**) are significantly different compared to the mock control (p-value < 0.03; wilcox test) have red p-values.

**Figure S2:**
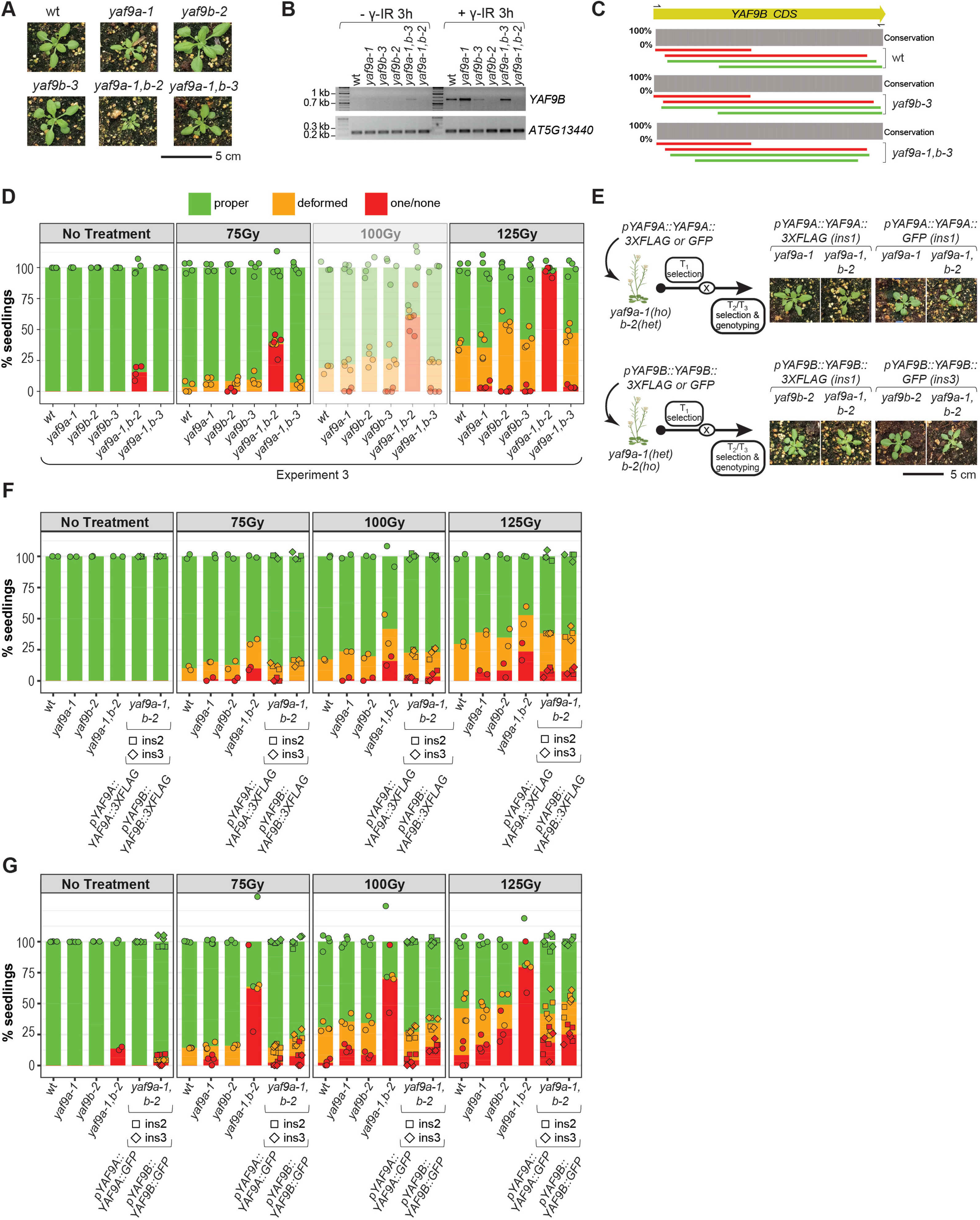
Characterization of *yaf9* mutants and complementing lines. (**A**) Images showing the phenotypes of 19-day-old seedlings in the indicated genotypes. (**B**) RT-PCR analysis of the full length *YAF9B* transcript (807 bp) in the indicated genotypes 3 hours after mock (- γ-IR) or γ-irradiation (+ γ-IR; 100Gy) treatments. Amplification of *AT5G13440* was used as a loading control. (**C**) Alignments showing the sequencing results of the full length *YAF9B* transcripts shown in **B** for the indicated genotypes. Forward and reverse sequences are shown in green and red, respectively, and the bar plots show the conservation compared to the *YAF9B* CDS. As the primers (black arrows) used to amplify *YAF9B* overlap with the start and end of the CDS, the sequences corresponding to these regions were independently verified from the mRNA-seq experiments for each genotype. (**D, F** and **G**) True leaf assays showing the percentage of 11-day-old seedlings categorized as having proper true leaves (green), deformed true leaves (orange) or one/none true leaves (red) after either no treatment or treatment with 75Gy, 100Gy, or 125Gy or gamma-irradiation. For each bar plot, the genotypes are labeled on the x-axis. On the y-axis, the color-coded dots represent the percentage of seedlings belonging to each category calculated based on sets of ∼30 seedlings that represent biological replicates (n=3 or 4). The stacked bars are the average of all the data points for each category. In D, the data for 100Gy is shown at a higher transparency to indicate it was also shown in Fig. 2B (experiment 3). In **F** and **G**, the data from two additional inserts of each tagged construct are shown as squares (ins2) and diamonds (ins3), with data for insert (ins1) shown in Fig. 2C. (**E**) Left: Schematics outlining the strategy used to generate 3XFLAG- or GFP-tagged YAF9A and YAF9B lines in the indicated mutant background. Right: Images showing the phenotypes of the resulting 19-day-old seedlings in the indicated genotypes.

**Figure S3:**
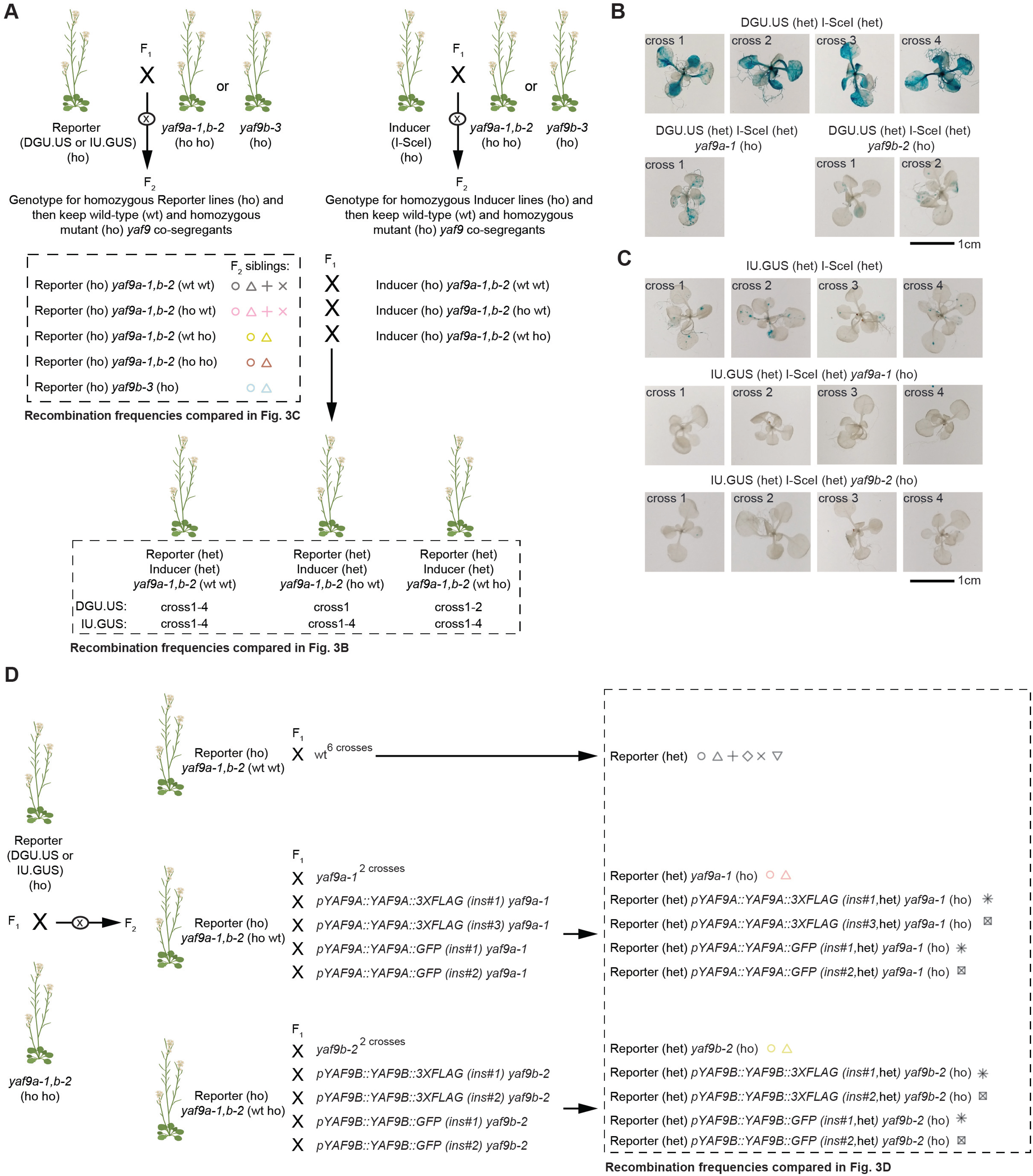
YAF9A and YAF9B are required for homologous recombination after targeted DSB damage. (**A**) Crossing scheme*s* to introduce the DGU.US and IU.GUS reporters or the I-SceI inducer (I-SceI) into yaf9 mutant backgrounds. The reporter lines (DGU.US or IU.GUS) as well as the I-SceI inducer line were independently crossed with either the *yaf9a-1,b-2* double mutant or the *yaf9b-3* single mutant. PCR-based genotyping in F_2_ plants allowed the identification *of* co-segregant plants either homozygous or wild-type for each *yaf9* allele. For the reporter crosses (Left), 2 to 4 independent F_2_ siblings for each genotype were identified, as represented by different symbols, and were used for the analyses presented in Fig. 3C. Reporter and inducer lines in the same genetic background (wt*, yaf9a-1, etc.*) were crossed together to generate plants heterozygous for both the reporter and the inducer construct but homozygous genetic background. For each genotype, multiple independent crosses (1-4) were conducted using unique parental lines to serve as biological replicates for the analysis shown in Fig. 3B. (**B** and **C**) Representative images of GUS stained 15-day-old seedlings showing recombination events at the DGU.US (**B**) and IU.GUS (**C**) reporters. For each set of images, the genotypes are indicated above and the cross numbers, which represent distinct biological replicates (as detailed in panel **A**), are indicated in the top-left corner of each image. Images from additional crosses are shown in Fig. 3B) (**D**) Crossing scheme to rescue HR-like defects in *yaf9* mutants using complementing lines. The reporter lines (DGU.US or IU.GUS) were independently crossed with the *yaf9a-1,b-2* double mutant. PCR-based genotyping in F_2_ plants allowed the identification of co-segregant plants that were homozygous for the reporter and either wt for both *yaf9* mutations (upper) or homozygous for just *yaf9a-1* (middle) or *yaf9b-2* (lower). These lines were then crossed with the indicated genotypes to generate the F_1_ seedlings used for Fig. 3D. In cases where multiple F_1_ crosses were conducted, the number of crosses are indicated and represented by different shapes in the genotyping summary. For the epitope-tagged YAF9 constructs, crosses were conducted using two independent lines (ins#) for each construct, which are also represented by different shapes in the genotyping summary.

**Figure S4:**
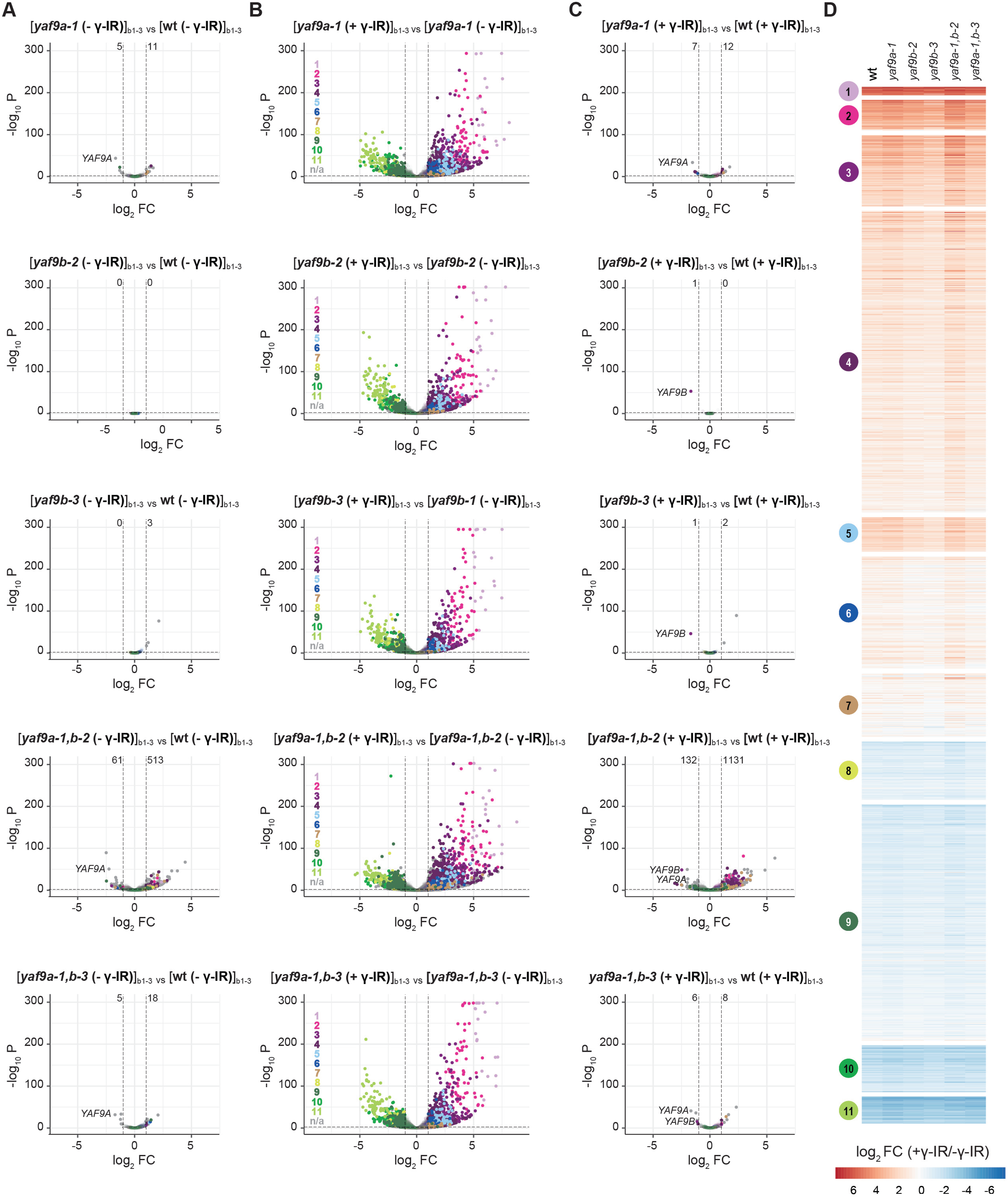
*yaf9* mutants show normal transcriptional responses upon exposure to DNA damage. (**A-C**) Volcano plots showing the changes in gene expression observed by comparing the genotypes and conditions indicated above each plot across three independent mRNA-seq experiments (b1-3). The x- and y-axes show the -log_10_ P and log_2_ FC values for each DEG and the dotted lines represent |log_2_ FC| ≥1 and p-value ≤0.01. In all cases, + γ-IR and - γ-IR correspond to 3h after gamma irradiation or mock treatments, respectively. DEGs present in the DREM model shown in Fig. 4A are color coded based on the gene groups in which they reside. DEGs not present in the DREM model are shown in gray. The dots corresponding to YAF9A and YAF9B are labeled and the additional DEGs are listed in **Table S2.** In **A** and **C**, the total number of up- and down-regulated genes are indicated to the left and right of the dotted lines marking the |log_2_ FC| ≥1. (**D**) Heatmaps showing the log_2_ FC in expression from the same comparisons shown in Fig. 4B and **S4B** for each of the genes present in the DREM model, separated into 11 groups. All heatmaps were generated using the indicated color scale.

**Figure S5:**
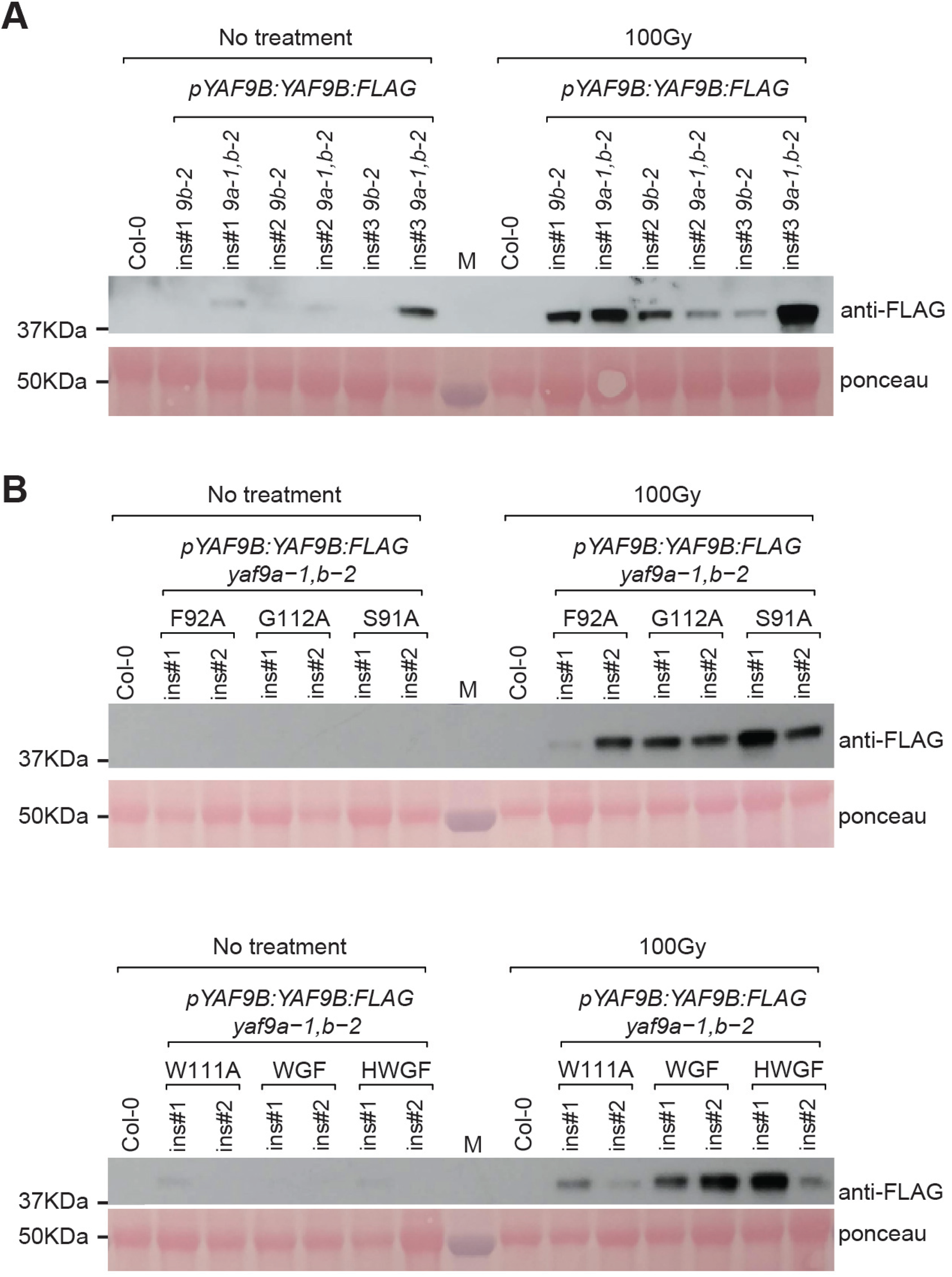
Wild-type and mutated versions of 3XFLAG-tagged YAF9B constructs are properly expressed in a damage-dependent manner. Anti-FLAG western blot detecting protein levels of 3XFLAG-tagged wt (**A**) and mutated (**B**) *YAF9B* 3h after gamma-irradiation (100Gy, + γ-IR) or no treatment (- γ-IR) from 2-3 independent transgenic plant lines, as well as a Col-0 control lacking a 3XFLAG transgene. Ponceau staining of the membrane is shown below as a loading control.

**Table S1: mRNA-seq sample descriptions and mapping statistics table.**

**Table S2: Gene expression matrix and DE gene tables.**

**Table S3: YAF9A and YAF9B Mass Spectrometry tables.**

**Table S4: Primers Table.**

**Table S5: Genetic constructs Tables.**

